# Molecular Mechanism of Hyperactivation Conferred by a Truncated TRPA1 Disease Mutant Suggests New Gating Insights

**DOI:** 10.1101/2022.01.05.475133

**Authors:** Avnika Bali, Samantha P. Schaefer, Isabelle Trier, Alice L. Zhang, Lilian Kabeche, Candice E. Paulsen

## Abstract

The wasabi receptor, TRPA1, is a non-selective homotetrameric cation channel expressed in primary sensory neurons of the pain pathway, where it is activated by diverse chemical irritants. A direct role for TRPA1 in human health has been highlighted by the discovery of genetic variants associated with severe pain disorders. One such TRPA1 mutant was identified in a father-son pair with cramp fasciculation syndrome (CFS) and neuronal hyperexcitability-hypersensitivity symptoms that may be caused by aberrant channel activity, though the mechanism of action for this mutant is unknown. Here, we show the CFS-associated R919* TRPA1 mutant is functionally inactive when expressed alone in heterologous cells, which is not surprising since it lacks the 201 C-terminal amino acids that house critical channel gating machinery including the pore-lining transmembrane helix. Interestingly, the R919* mutant confers enhanced agonist sensitivity when co-expressed with wild type (WT) TRPA1. This channel hyperactivation mechanism is conserved in distant TRPA1 species orthologues and can be recapitulated in the capsaicin receptor, TRPV1. Using a combination of ratiometric calcium imaging, immunostaining, surface biotinylation, pulldown assays, fluorescence size exclusion chromatography, and proximity biotinylation assays, we show that the R919* mutant co-assembles with WT subunits into heteromeric channels. Within these heteromers, we postulate that R919* TRPA1 subunits contribute to hyperactivation by lowering energetic barriers to channel activation contributed by the missing regions. Additionally, we show heteromer activation can originate from the R919* TRPA1 subunits, which suggests an unexpected role for the ankyrin repeat and coiled coil domains in concerted channel gating. Our results demonstrate the R919* TRPA1 mutant confers gain-of-function thereby expanding the physiological impact of nonsense mutations, reveals a novel and genetically tractable mechanism for selective channel sensitization that may be broadly applicable to other receptors, and uncovers new gating insights that may explain the molecular mechanism of temperature sensing by some TRPA1 orthologues.

## INTRODUCTION

The wasabi receptor, TRPA1 (<u>T</u>ransient <u>R</u>eceptor <u>P</u>otential <u>A</u>nkyrin subtype <u>1</u>), is a calcium-permeable non-selective homotetrameric cation channel expressed in a subset of primary sensory neurons of the dorsal root, trigeminal, and nodose ganglia where it plays a role in many pain-associated sensory disorders including itch, airway diseases, and inflammation^1–6^. The TRPA1 chemosensor is activated by a chemically diverse panel of environmental and endogenous toxins that fall into two broad categories: electrophiles and non-electrophiles. Electrophile agonists such as Allyl Isothiocyanate (AITC) and Cinnamaldehyde activate the channel by covalently modifying key conserved cysteine residues in the membrane proximal cytoplasmic N-terminus^7–10^. Non-electrophile agonists such as Carvacrol bind at distinct sites in the transmembrane domain to promote channel activation^11–14^. Recent cryo-electron microscopy TRPA1 structures captured in the closed and electrophile agonist activated states reveal large, allosterically coupled conformational changes in the transmembrane domain and membrane-proximal cytoplasmic N- and C-termini, referred to as the allosteric nexus, highlighting key roles for these regions in channel gating^15–17^.

The TRPA1 transmembrane domain and allosteric nexus account for only 35% of the total protein sequence (390 of 1119 amino acids) and little is known about the functional impact of the remaining cytoplasmic domains. Situated below the allosteric nexus, TRPA1 houses a large N-terminal ankyrin repeat domain (ARD), in which the five membrane proximal ARs form a cage around a C-terminal coiled coil that coordinates a requisite polyanion cofactor (Fig. 1a-c)^15,18^. Interestingly, certain TRPA1 species orthologues can be activated by temperature, though the molecular mechanism of temperature sensing is unclear^19^. Studies with TRPA1 species chimeras and point mutations have shown thermosensation can be conferred or tuned through the ARD^20,21^. Whether or how the ARD can communicate to the channel pore remains unknown.

**Figure 1.**
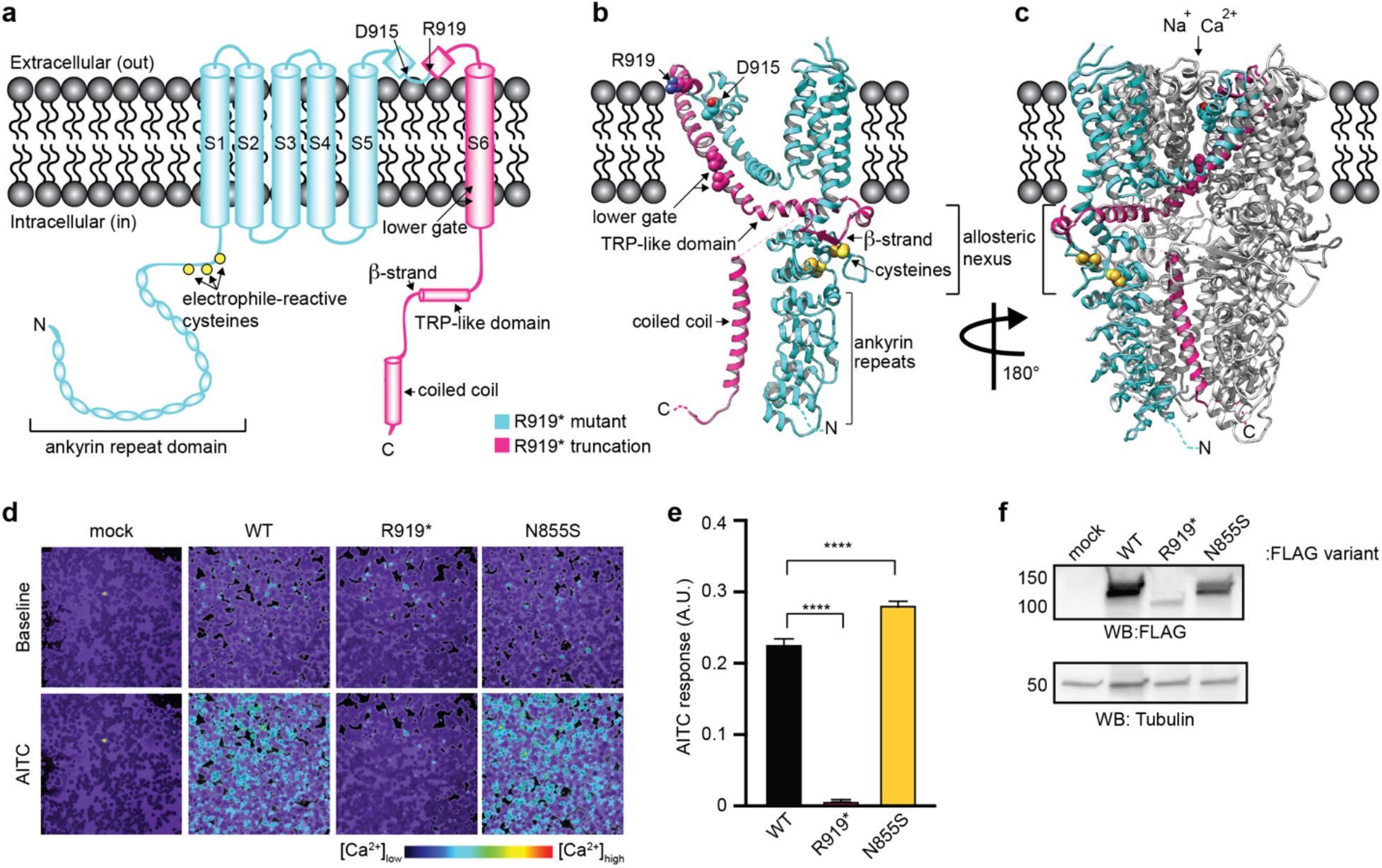
The R919* mutant is a nonfunctional TRPA1 natural variant. (a) Cartoon schematic of a full-length hTRPA1 monomeric subunit, with relevant structural features denoted. Regions retained in R919* hTRPA1 are indicated in teal. Regions truncated in R919* hTRPA1 are indicated in pink. (b-c) Ribbon diagrams of WT hTRPA1 atomic model for residues K446-T1078 from a single subunit (b) and the homotetrameric channel (c). Color scheme and relevant structural features denoted as in (a). In (c), only one subunit is colored for clarity. Allosteric nexus additionally indicated with brackets. Models built with the human TRPA1 Cryo-EM structure (PDB: 6V9W) in UCSF Chimera. (d) Ratiometric calcium imaging of HEK293T cells transiently transfected with empty vector (mock), WT hTRPA1, R919* hTRPA1, or N855S hTRPA1. Cells were stimulated with AITC (1θ0 μM). Images are representative of three independent experiments. (e) Quantification of AITC-evoked change in Fura-2 ratio (arbitrary units, AU). Data represent mean ± SEM. n = 4 independent experiments, n ≥ 90 cells per transfection condition per experiment. ****p<0.0001, one-way ANOVA with Bonferroni’s *post hoc* analysis. (f) Western blot of lysates from transiently transfected HEK293T cells expressing 3xFLAG-tagged hTRPA1 variants, probed using HRP-conjugated anti-FLAG antibody. Tubulin was the loading control. Results were verified in four independent experiments.

Ion channel mutations with associated pathological conditions, known as channelopathies, provide direct evidence a channel is relevant to human health, and can illuminate molecular mechanisms and protein domains that govern normal and aberrant channel function^22^. To date, two human TRPA1 disease mutations associated with congenital pain disorders have been reported^23,24^, which illustrates a direct role for TRPA1 in pain signaling, marking it as a promising therapeutic target. The first characterized TRPA1 channelopathy, which was discovered in patients with a rare, autosomal dominant Familial Episodic Pain Syndrome (FEPS), introduced a missense mutation at N855^23^. The N855S TRPA1 mutant forms functional channels when expressed alone in heterologous cells characterized by increased inward current at resting membrane potentials with no change to agonist sensitivity. The location of the N855S mutation in a short linker that physically couples two transmembrane helical bundles highlighted a key role for this region in channel gating years before high-resolution TRPA1 structures were available.

The second reported TRPA1 channelopathy has not yet been characterized and a mechanism for how it may alter channel function is unknown. This mutant was discovered in a father-son pair with Cramp Fasciculation Syndrome (CFS) and other neuronal hyperexcitability-hypersensitivity symptoms including cold hyperalgesia, chronic itch, and asthma that are consistent with aberrant TRPA1 function^24^. The CFS *TRPA1* variant introduces a nonsense mutation at R919 located at the start of the second pore helix, and the resulting R919* TRPA1 protein lacks the final 201 amino acids, including critical elements involved in gating such as the pore-lining transmembrane helix (S6) and components of the allosteric nexus (Fig. 1a-c). Given the degree of truncation of the R919* TRPA1 mutant, a haploinsufficiency or loss-of-function phenotype might be expected akin to *TRPA1* heterozygous and homozygous knockout animals, respectively^25,26^. However, the possible mechanisms by which the R919* TRPA1 mutant may underlie hyperexcitability-hypersensitivity symptoms are numerous, given that both gain-of-function and loss-of-function mutations in voltage-gated sodium channels have been associated with pronounced hyperexcitability phenotypes in human patients^27,28^. In this study, we sought to elucidate whether and how the R919* TRPA1 mutant influences channel activity that could explain the observed congenital pain phenotype in patients.

## RESULTS

### Functional Characterization of the R919* TRPA1 mutant

Whole-exome sequencing and co-segregation analysis of a father-son pair with CFS and other hypersensitivity-hyperexcitability symptoms identified a novel TRPA1 gene variant consisting of a C to T transition (c.2755C>T)^24^. When mapped onto the wild type (WT) human TRPA1 monomer structure, the R919* mutant truncates the last 201 amino acids, or 18% of the protein, removing many important structural and regulatory features including the second pore helix, the ion conduction pathway-lining S6 transmembrane helix and the entire cytoplasmic C-terminus (Fig. 1a-c). If R919* TRPA1 contributes to the hypersensitivity-hyperexcitability symptoms exhibited by patients carrying this variant, the R919* mutant might form channels with gain-of-function properties akin to the previously reported FEPS-associated TRPA1 mutant, N855S^23^. Channelopathies are commonly studied in isolation in heterologous systems to reveal alterations to channel biophysical properties^23,29–32^. Thus, human R919* TRPA1 was expressed in HEK293T cells, which do not express TRPA1, and channel activity was assayed by Fura-2 ratiometric calcium imaging (Fig. 1d and e, Extended Data Fig. 1a). While robust activation of WT TRPA1 was observed by the electrophile agonist AITC (Fig. 1d and e) and the non-electrophile agonist Carvacrol (Extended Data Fig. 1a), and enhanced activation of N855S TRPA1 was observed with both agonists consistent with its reported gain-of-function properties, cells expressing the R919* TRPA1 mutant revealed no activity with either agonist (Fig. 1d and e, Extended Data Fig. 1a).

Immunoblot analysis confirmed R919* mutant protein was produced in HEK293T cells, albeit at significantly lower levels than WT or N855S TRPA1 (Fig. 1f). Nonetheless, WT TRPA1 still formed functional channels when expressed at a comparable amount (Extended Data Fig. 1b and C). These results suggest that the lack of detectable activity for R919* TRPA1 is due to its inability to produce functional calcium permeable ion channels, which is unsurprising given the loss of the many key structural and functional domains from this mutant (Fig. 1a-c).

### Functional Characterization of WT and R919* TRPA1 Co-Expression

Both R919* and N855S TRPA1 mutants were discovered in heterozygous individuals, where WT and mutant TRPA1 protein could be co-produced^23,24^. This raised the possibility that the R919* mutant mediates its effects by altering co-expressed WT TRPA1 properties. To test whether the R919* mutant affected channel activity in the presence of WT TRPA1, ratiometric calcium imaging was performed on HEK293T cells co-expressing both variants (Fig. 2). Cells co-expressing WT and R919* TRPA1 exhibited a robust increase in calcium influx compared to cells expressing WT TRPA1 alone, especially in response to sub-saturating concentrations of AITC (Fig. 2a and b) and Carvacrol (Extended Data Fig. 2a). There was also a marked enhancement in the time to maximal response at sub-saturating AITC concentrations (Fig. 2b). Together, these observations suggest that co-expression of WT TRPA1 with the R919* mutant hyperactivates channels. These hyperactive channels were inhibited by the canonical TRPA1 antagonists A-967079, HC-030031, and ruthenium red, consistent with TRPA1 activity (Fig. 2c, Extended Data Fig. 3).

**Figure 2.**
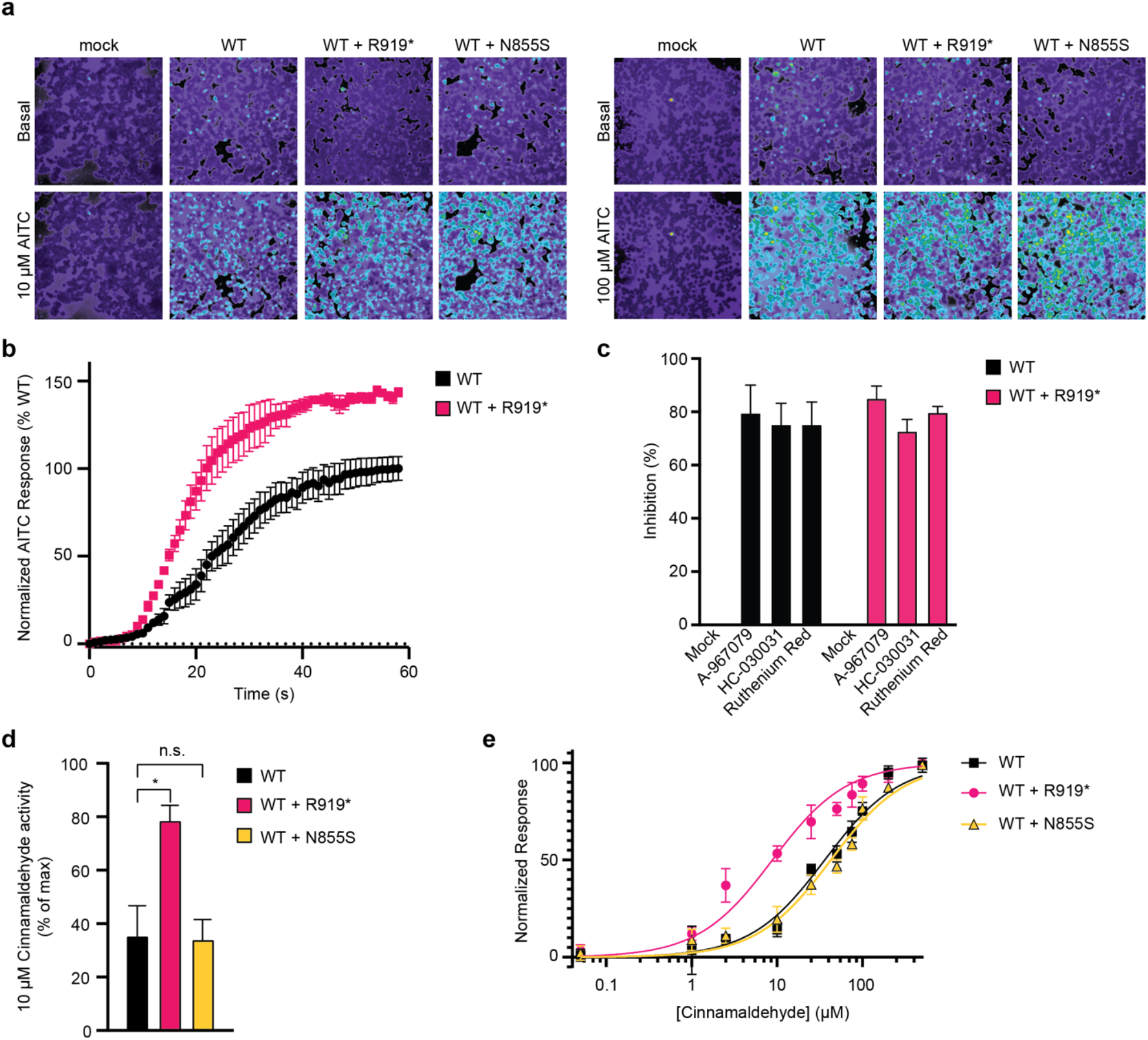
The R919* mutant confers hyperactivity when co-expressed with WT TRPA1. (a) Ratiometric calcium imaging of HEK293T cells transiently transfected with empty vector (mock), WT hTRPA1, WT and R919* hTRPA1, or WT and N855S hTRPA1. Cells were stimulated with 10 μM (left) or 100 μM AITC (right). Images are representatives from three independent experiments. (b) Time-dependent ratiometric calcium imaging responses evoked by 10 μM AITC normalized to maximum response elicited from WT hTRPA1-expressing cells. Data represent mean ± SEM. n = 3 independent experiments, n ≥ 90 cells per condition per experiment. (c) Quantification of antagonist-mediated inhibition of 100 μM AITC-evoked changes in Fura-2 ratio in HEK293T cells transiently transfected with WT hTRPA1 or WT and R919* hTRPA1. Cells were pre-treated with Ringer’s solution (mock), A-967079 (10 μM), HC-030031 (30 μM), or Ruthenium Red (10 μM). Data represent mean ± SEM of percentage of inhibition of the AITC-evoked maximum response for each transfection type. n = 4 independent experiments, n ≥ 90 cells per condition per experiment. (d) Quantification of 10 μM Cinnamaldehyde-evoked change in Fura-2 ratio relative to maximum response of each expression condition at 100 μM Cinnamaldehyde. Data represent mean ± SEM. *p<0.05, n.s. not significant. n = 3 independent experiments, n ≥ 90 cells per condition per experiment, one-way ANOVA with Bonferroni’s *post hoc* analysis. (e) Dose-response curve of cinnamaldehyde-evoked calcium responses for HEK293T cells transiently transfected with WT hTRPA1, WT and R919* hTRPA1, or WT and N855S hTRPA1. Calcium responses normalized to maximum calcium response to 500 μM Cinnamaldehyde. Traces represent the average ± SEM of normalized calcium responses from 3 independent experiments, n = 30 cells per agonist concentration per experiment. Data were fit to a non-linear regression. EC_50_ (95% CI) values are 37.7 μM for WT hTRPA1 (95% CI, 30.5-45.8 μM), 7.8 μM for WT and R919* hTRPA1 (95% CI, 6.1-9.9 μM), and 43.1 μM for WT and N855S hTRPA1 (95% CI, 33.1-54.5 μM). Cinnamaldehyde EC_50_ reduction for WT and R919* hTRPA1 co-expression is statistically significant (p<0.0001, Extra sum-of-squares F test).

Enhanced channel activity specifically at sub-saturating AITC concentration suggests that co-expression of WT and R919* TRPA1 affects agonist sensitivity. This is distinct from cells coexpressing WT and N855S TRPA1, which displayed a clear enhancement of calcium influx at both sub-saturating and saturating AITC concentrations (Fig. 2a). Despite an increase in calcium influx at each agonist concentration, the N855S mutant agonist sensitivity was previously shown to be no different from WT TRPA1 with the electrophile agonist Cinnamaldehyde^23^. To assess agonist sensitivity, channel activity at a sub-saturating Cinnamaldehyde concentration (10 μM) was normalized to maximum channel activity at a saturating Cinnamaldehyde concentration (100 μM). Consistent with previously published work, no change from WT TRPA1 Cinnamaldehyde sensitivity was detected when co-expressed with the N855S mutant (Fig. 2d)^23^. In contrast, a significant increase in Cinnamaldehyde sensitivity from cells co-expressing WT and R919* TRPA1 was observed (Fig. 2d). Cinnamaldehyde sensitivity was quantified by determining the half maximum effective concentration (EC_50_) from cells expressing WT TRPA1 alone or with the N855S or R919* mutants. While no significant change in the Cinnamaldehyde EC_50_ was observed when WT TRPA1 was co-expressed with the N855S mutant (37.7 μM for WT TRPA1 versus 43.1 μM for WT and N855S TRPA1), a 4.8-fold increase in Cinnamaldehyde sensitivity (EC_50_ = 7.8 μM) was observed with R919* TRPA1 co-expression (Fig. 2e)^23^. Significantly enhanced agonist sensitivity was also observed with Carvacrol, indicating the R919* mutant mediates channel hypersensitivity independent of agonist type (Extended Data Fig. 2b and c).

Collectively, these results suggest that the R919* mutant confers enhanced activation kinetics and agonist sensitivity when co-expressed with WT TRPA1, consistent with mutant-mediated sensitization and hyperactivation of TRPA1 channels (Fig. 2). In contrast, the N855S mutant has no effect on agonist sensitivity. Thus, the mechanism of channel hyperactivity conferred by the R919* mutant is likely distinct from N855S TRPA1.

### Effect of the R919* Mutant on WT TRPA1 Expression, Localization, and General Cell Stress

Channels can become hyperactive through altered protein stability, enhanced plasma membrane trafficking or via post-translational channel modifications triggered by general cell stress^33–43^. Immunoblot analysis indicated that expression of R919* had no effect on the level of WT TRPA1 (Extended Data Fig. 4a and b). Identical results were obtained from cells used in parallel for ratiometric calcium imaging (Extended Data Fig. 4c). Similarly, individual or co-expression of WT and R919* TRPA1 had no overt effects on cell viability as assessed by Trypan blue exclusion (Extended Data Fig. 4d). It is possible R919* TRPA1 causes cell stress capable of hypersensitizing ion channels that was not detected by Trypan blue exclusion. Thus, R919* TRPA1 was co-expressed with the related capsaicin receptor, TRPV1 and assayed for its effect on human TRPV1 expression, activity, or agonist sensitivity. Akin to WT TRPA1, the R919* mutant had no effect on TRPV1 expression (Extended Data Fig. 4a and b). Moreover, coexpression with R919* TRPA1 conferred no change in capsaicin evoked TRPV1 calcium influx or agonist sensitivity (Extended Data Fig. 5). Finally, surface biotinylation assays revealed that coexpression of R919* had no effect on the amount of WT TRPA1 at the plasma membrane (Fig. 3a and b). Together, these data indicate that the R919* mutant does not sensitize channels via changes in WT TRPA1 expression, altered plasma membrane localization, or general induction of cell stress.

**Figure 3.**
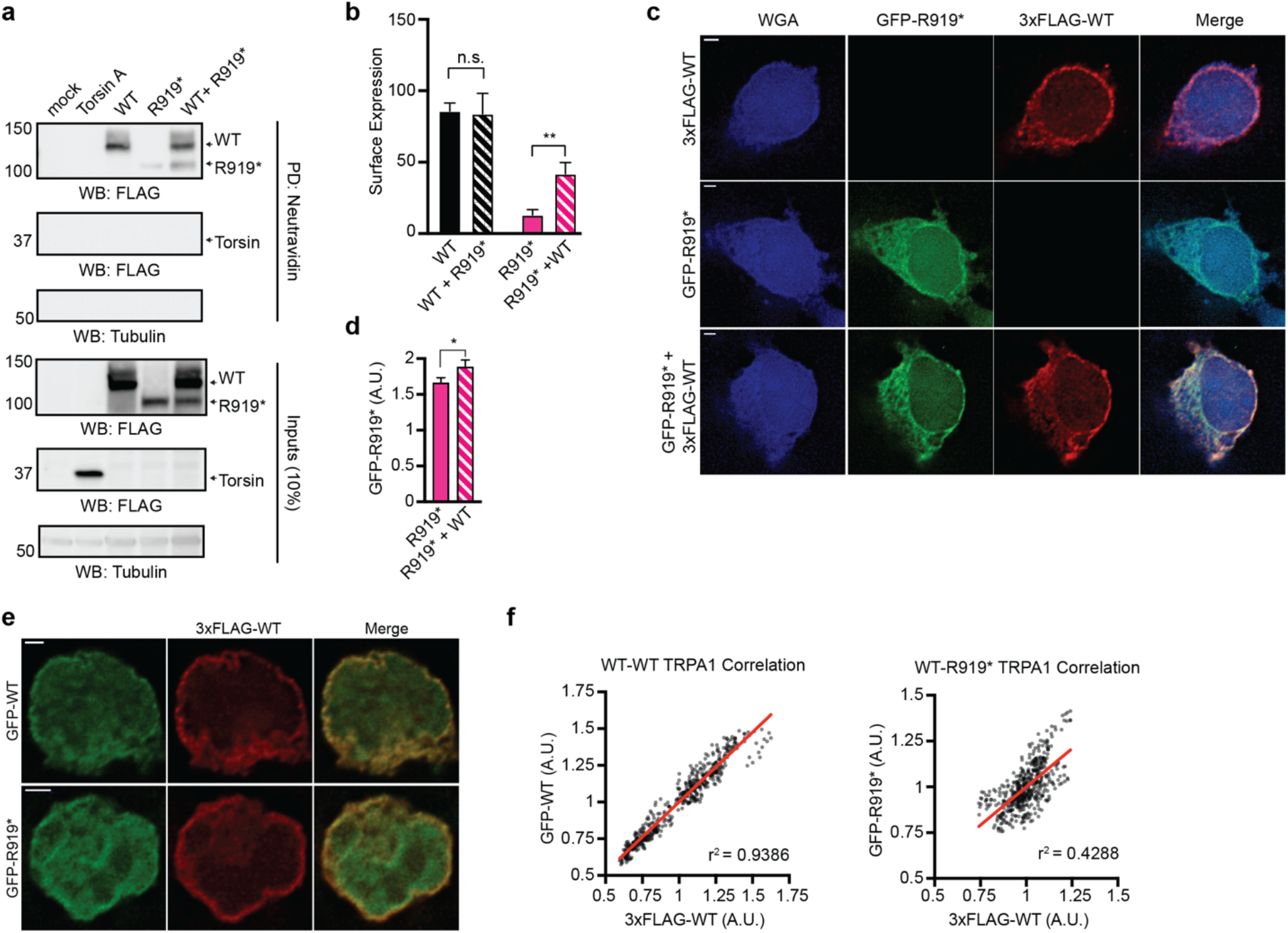
WT and R919* TRPA1 co-localize at the plasma membrane in cells. (a) Immunoblotting analysis of 3xFLAG-WT hTRPA1, 3xFLAG-R919* hTRPA1, or FLAG-Torsin A protein expression in biotin-labeled plasma membranes from transiently transfected HEK293T cells. Biotinylated proteins were precipitated by Neutravidin resin pulldown and probed using HRP-conjugated anti-FLAG antibody. Tubulin from whole cell lysates (10%, inputs) was the loading control. Torsin A and Tubulin were the negative controls for plasma membrane localization. (b) Quantitative analysis of the localization of 3xFLAG-WT hTRPA1 or 3xFLAG-R919* hTRPA1 proteins in the plasma membrane relative to Tubulin. WT+R919* and R919*+WT indicate the expression of WT or R919* hTRPA1 in the plasma membrane of co-transfected cells, respectively and are represented by striped bars. Data represent mean ± SEM. **p<0.01, n.s. not significant. n=6, Student’s t-test. (c) Representative deconvolved immunofluorescence images of HEK293T cells transiently transfected with GFP-R919* hTRPA1, 3xFLAG-WT hTRPA1, or GFP-R919* hTRPA1 and 3xFLAG-WT hTRPA1. Cells were stained with anti-GFP (green) and anti-FLAG (red) antibodies. Plasma membrane was labeled with wheat germ agglutinin (blue). Scale bar indicates 2 μm. (d) Quantification of GFP-R919* hTRPA1 fluorescence intensity in the plasma membrane of HEK293T cells relative to the cell interior. Values were obtained using line-scans of raw images. Data represent mean ± SEM. *p<0.05, n=30 cells per condition, Student’s t-test. (e) Representative deconvolved immunofluorescence images of HEK293T cells co-expressing 3xFLAG-WT hTRPA1 with GFP-WT hTRPA1 or GFP-R919* hTRPA1. Cells were stained with anti-GFP (green) and anti-FLAG (red) antibodies. Scale bar indicates 2 μm. (f) Scatter plot of 3xFLAG-WT hTRPA1 and GFP-WT hTRPA1 (left) or GFP-R919* hTRPA1 (right) fluorescence intensity at the plasma membrane of HEK293T cells. Pearson’s correlation coefficients (r) were determined using raw images and the coefficient of determination (r^2^) is depicted in the lower right corner of each plot. A line of best fit is shown in red. p<0.0001, n>450 pixels in 1 cell per condition.

### Effect of WT TRPA1 on R919* Mutant Expression and Localization

Some but not all ion channel truncation mutants result in defects in plasma membrane trafficking^38,42,44–49^. Surface biotinylation assays revealed a small plasma membrane population of the R919* mutant, which increased significantly when co-expressed with WT TRPA1 (Fig. 3a and b). Torsin A, an endoplasmic reticulum and perinuclear space-resident AAA+ ATPase, was used as an internal negative control to ensure the R919* mutant labeling was attributed to true plasma membrane localization and not to probe internalization during such assays (Fig. 3a)^50^. We further investigated the R919* mutant plasma membrane localization using immunofluorescence imaging. Deconvolved images of HEK293T cells expressing GFP-tagged R919* TRPA1 or 3xFLAG-tagged WT TRPA1 revealed robust WT TRPA1 localization at the cell surface, while the R919* mutant displayed aberrant localization spreading more diffusely throughout the cytoplasm with minimal plasma membrane localization (Fig. 3c). When co-expressed, 3xFLAG-tagged WT and GFP-tagged R919* TRPA1 both localized at the plasma membrane (Fig. 3c). Cross-sectional line scans were performed on raw images of immuno-stained cells, revealing a statistically significant increase in R919* mutant at the cell edge when co-expressed with WT TRPA1 (Fig. 3d and Extended Data Fig. 6a and b). Finally, co-expression with WT TRPA1 did not increase the R919* mutant protein expression, suggesting enhanced plasma membrane localization is not simply due to stabilization of R919* TRPA1 protein (Fig. 3a, Extended Data Fig. 4a and b). Together, these data suggest that co-expression of WT and R919* TRPA1 influences the trafficking behavior of the R919* mutant.

### Co-Localization and Physical Interaction of WT and R919* TRPA1 in Cells

A direct interaction between WT and R919* TRPA1 would explain the ability of WT TRPA1 to enhance the surface localization of the R919* mutant (Fig. 3a-d), and this interaction could structurally impact WT TRPA1 to confer the observed channel sensitization (Fig. 2). Indeed, deconvolved immunofluorescence images of HEK293T cells co-expressing GFP-tagged R919* TRPA1 and 3xFLAG-tagged WT TRPA1 exhibit apparent co-localization of WT and R919* TRPA1 at the plasma membrane (Fig. 3c and e). Protein co-localization can be further analyzed by line scan analysis of plasma membrane segments on raw images of immuno-stained cells^51,52^. Such analysis on HEK293T cells co-expressing GFP-tagged WT TRPA1 and 3xFLAG-tagged WT or R919* TRPA1 revealed a strong positive correlation between WT subunits that is consistent with known channel homotetramerization, and a positive, albeit weaker correlation between WT and R919* TRPA1 signal at the cell surface (Fig. 3f and Extended Data Fig. 7c-e).

Since WT and R919* TRPA1 co-localize in cells, it is possible these proteins physically interact. To characterize such an interaction, pulldown assays were conducted on lysates from cells co-expressing differentially tagged variants of WT or R919* TRPA1. These pulldown assays revealed a robust interaction between MBP-tagged WT TRPA1 and 3xFLAG-tagged WT and R919* TRPA1, but not with 3xFLAG-tagged Kv1.2/2.1 (Kv), a voltage-gated potassium channel (Fig. 4a). Similarly, MBP-tagged R919* TRPA1 was able to efficiently pulldown 3xFLAG-tagged WT and R919* TRPA1, with no interaction observed with 3xFLAG-tagged Kv (Fig. 4a).

**Figure 4.**
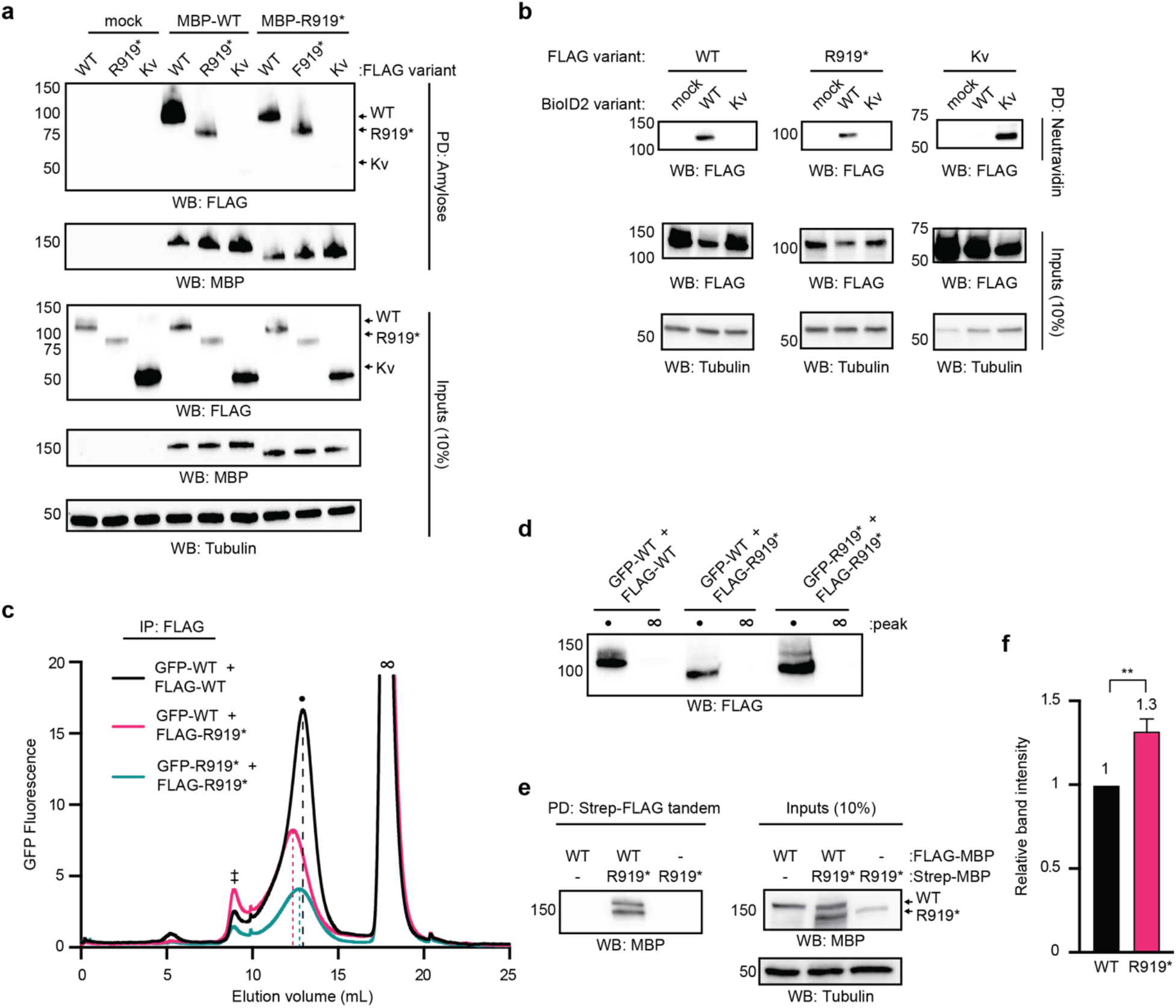
WT and R919* TRPA1 associate in cells to form complexes of similar size to homotetrameric WT channels. (a) Immunoblotting analysis of 3xFLAG-WT hTRPA1, 3xFLAG-R919* hTRPA1, or 3xFLAG-Kv1.2/2.1 protein expression after amylose pulldown from lysates of HEK293T cells transiently co-transfected with empty vector (mock), MBP-WT hTRPA1, or MBP-R919* hTRPA1. Samples were probed using HRP-conjugated anti-FLAG antibody. MBP-WT hTRPA1 and MBP-R919* hTRPA1 were probed using anti-MBP antibody. Tubulin from whole cell lysates (10%, inputs) was the loading control. Results were verified in 3 independent trials. (b) Immunoblotting analysis of biotinylated 3xFLAG-WT hTRPA1,3xFLAG-R919* hTRPA1, or 3xFLAG-Kv1.2/2.1 co-expressed in HEK293T cells with empty vector (mock), BioID2-WT hTRPA1, or BioID2-Kv1.2/2.1. Biotinylated proteins were precipitated by Neutravidin resin pulldown and probed using HRP-conjugated anti-FLAG antibody. Tubulin from whole cell lysates (10%, inputs) was the loading control. Results were verified in 3 independent trials. (c) FSEC chromatograms from GFP-WT hTRPA1 and GFP-R919* hTRPA1 transiently co-transfected in HEK293T cells with 3xFLAG-WT hTRPA1 or 3xFLAG-R919* hTRPA1. FLAG immunoprecipitated eluents were analyzed by FSEC. Chromatograms reveal elution profiles of co-purified GFP-WT hTRPA1 or GFP-R919* hTRPA1 complexes. Peaks corresponding to void (‡), tetrameric WT hTRPA1 channels (•) and free GFP (∞) are indicated. Dashed lines denote the center elution volume of each co-purified complex. Results were verified in 3 independent trials. (d) Immunoblotting analysis of 3xFLAG-WT or R919* hTRPA1 from indicated peak fractions from (C) probed using HRP-conjugated anti-FLAG antibody. (e) Immunoblotting analysis of tandem-purified WT-R919* hTRPA1 complexes. FLAG-MBP-WT hTRPA1 and Strep-MBP-R919* hTRPA1 were transiently transfected separately or together in HEK293T cells. Lysates were tandem purified for Strep-then FLAG-tagged proteins. MBP-tagged proteins of tandem purification eluents were probed using anti-MBP antibody. Tubulin from whole cell lysates (10%, inputs) was the loading control. (f) Quantitative analysis of FLAG-MBP-WT hTRPA1 and Strep-MBP-R919* hTRPA1 from tandem purifications in (E). Band intensity was normalized to WT hTRPA1. Data represent mean ± SEM. Means are indicated above the bars. **p<0.01, n=3, Student’s t-test.

A proximity biotinylation ligation assay was used to further investigate whether WT and R919* TRPA1 directly interact in intact cells^53^. BioID2-fused WT TRPA1 or Kv were co-expressed with 3xFLAG-tagged WT TRPA1, R919* TRPA1, or Kv. Biotinylated 3xFLAG-tagged variants were detected by immunoblot analysis of Neutravidin eluates. These experiments revealed that 3xFLAG-tagged WT and R919* TRPA1 were biotinylated by BioID2-fused WT TRPA1, but not by BioID2-fused Kv (Fig. 4b). Additionally, 3xFLAG-tagged Kv was biotinylated by BioID2-fused Kv, but not by BioID2-fused WT TRPA1 (Fig. 4b). Collectively, these results indicate that WT and R919* TRPA1 co-localize at the plasma membrane and engage in a close-range physical interaction in cells.

### Biochemical Characterization of Isolated WT and R919* TRPA1 Complexes

The proximity biotinylation results reveal that WT and R919* TRPA1 subunits are within a range of ~10 nm, approximately the width of a single TRPA1 channel^15^. This labeling radius is consistent with proximity of adjacent WT and R919* TRPA1 homotetramers or with heteromeric complexes formed by co-assembly of WT and R919* TRPA1 subunits. While TRPA1 is the only member of the TRPA sub-family of mammalian TRP ion channels, heteromerization within the multi-member TRPV, TRPC, TRPP, and TRPML sub-families is well characterized^54–64^, raising the possibility that WT and R919* TRPA1 subunits could heteromerize in a similar manner. Indeed, such heteromerization of WT subunits with alternative splice variants or single nucleotide polymorphisms has been proposed for TRPV1 and TRPA1, respectively^65,66^.

Fluorescence size exclusion chromatography (FSEC) was used to differentiate between association of adjacent homotetramers versus assembly of heteromeric complexes as these assemblies are of very different sizes *(e.g.,* ~1000 KD versus ~500 KD, respectively) and would yield distinct elution profiles^67^. GFP- or 3xFLAG-tagged WT or R919* TRPA1 were co-expressed in HEK93T cells in specific combinations, complexes were isolated by anti-FLAG immunoprecipitation, and eluates were analyzed by FSEC to monitor the associated GFP-tagged subunits. The elution profile from cells co-expressing GFP- or 3xFLAG-tagged WT TRPA1 variants revealed a monodisperse peak consistent with a WT TRPA1 homotetramer, as previously reported (Fig. 4c, black trace)^15^. These complexes were of similar size to homotetrameric GFP-WT TRPA1 channels suggesting the heterogeneity of a large GFP tag and a small 3xFLAG tag had minimal impact on the elution profile (Extended Data Fig. 7a). Complexes isolated from cells co-expressing GFP- or 3xFLAG-tagged R919* TRPA1 revealed a monodisperse peak that eluted slightly before WT TRPA1 homotetramers (Fig. 4c, teal trace). Interestingly, complexes isolated from cells co-expressing GFP-tagged WT TRPA1 and 3xFLAG-tagged R919* TRPA1 revealed a monodisperse peak that elutes slightly earlier than WT or R919* homotetramers (Fig. 4c, pink trace). Immunoblot analysis of FSEC fractions from the TRPA1 (•) and free GFP (∞) peaks confirmed the presence of 3xFLAG-tagged variants in the TRPA1 peaks (Fig. 4d).

Size exclusion chromatography elution profiles are affected by both protein size and shape thus, to further interrogate the oligomer state of WT-R919* complexes, these assemblies were isolated by tandem purification of lysates from HEK293T cells expressing dual-tagged FLAG-MBP- or Strep-MBP-WT or R919* TRPA1 variants and analyzed by Blue Native PAGE. The results of such experiments further confirm WT-R919* complexes resemble WT TRPA1 homotetramers in size (Extended Data Fig. 7b). This is distinct from similar tandem-purified complexes of WT mouse TRPA1 and the TRPA1b alternative splice variant, which ran markedly larger than WT homotetrameric complexes on Blue Native PAGE^41^. Notably, CFS homomers were not resolved after tandem purification by this method, perhaps suggesting their relative instability and/or revealing transient interactions between R919* TRPA1 subunits (Extended Data Fig. 7b).

There are three possible stoichiometries for the assembly of WT and R919* TRPA1 subunits into heterotetramers: 3:1, 2:2, or 1:3 WT:R919* TRPA1 subunits. To estimate subunit stoichiometry in WT-R919* TRPA1 heteromeric complexes, these assemblies were isolated by Strep-FLAG tandem purification as above. Immunoblot analysis with a common MBP-tag readout confirmed the isolation of such complexes only from cells co-expressing WT and R919* TRPA1 variants (Fig. 4e). The common MBP-tag readout additionally facilitated quantification of WT and R919* TRPA1 band intensities of these isolated assemblies, which yielded an average WT:R919* TRPA1 subunit ratio of 1:1.3 (Fig. 4f). This subunit ratio is inconsistent with a single WT-R919* TRPA1 heteromer composition. Together, these data suggest that WT and R919* TRPA1 subunits co-assemble into stable, tetrameric channel-sized complexes of heterogeneous subunit stoichiometries.

### Assessment of R919* TRPA1 Subunit Contribution to Functional Channels

Co-assembly of R919* and WT TRPA1 subunits into heteromeric channel-sized assemblies raises the possibility that these heteromeric complexes form functional channels that are a direct cause of the observed hyperactivity. Electrophile agonist complementation assays were performed to investigate whether R919* TRPA1 subunits contribute to functional channels. AITC activates TRPA1 by covalent modification of three key conserved cysteine residues in the cytoplasmic N-terminus *(e.g.,* C621, C641, and C665) that are present in both WT and R919* TRPA1 subunits (Fig. 1a and 5a)^8,10^. Mutation of these three cysteines to serine on WT TRPA1 yields a full-length construct (denoted 3CtoS FL) with greatly reduced AITC sensitivity, while preserving activation by the non-electrophile agonist Carvacrol (Fig. 5a and b, Extended Data Fig. 8a and b). Thus, AITC sensitivity of 3CtoS FL TRPA1 can be quantified by normalizing AITC-evoked calcium influx to Carvacrol-evoked activity (Fig. 5a). To test whether the R919* mutant subunits contribute to functional channels, the R919* variant was co-expressed with 3CtoS FL TRPA1 in cells and assayed for AITC-evoked activity compared to 3CtoS FL TRPA1 alone. Such experiments revealed that 3CtoS FL TRPA1 AITC sensitivity could be significantly restored by coexpression with the R919* mutant, suggesting that R919* subunits can complement the loss of reactive residues (Fig. 5b, compare teal and dark purple bars, Extended Data Fig. 8a and b). This functional rescue was contingent on the R919* mutant subunits themselves containing the three cysteine residues since co-expression of 3CtoS FL TRPA1 with the 3CtoS R919* mutant failed to enhance AITC-evoked calcium influx (Fig. 5B, lavender bar, Extended Data Fig. 8a and b). Together, these results support direct co-assembly of WT and R919* TRPA1 subunits into functional heteromeric channels with concerted activation among subunits as is typical of homotetrameric TRPA1 channels^68^. Moreover, these results demonstrate electrophile-mediated channel activation can originate from the R919* mutant subunit(s).

**Figure 5.**
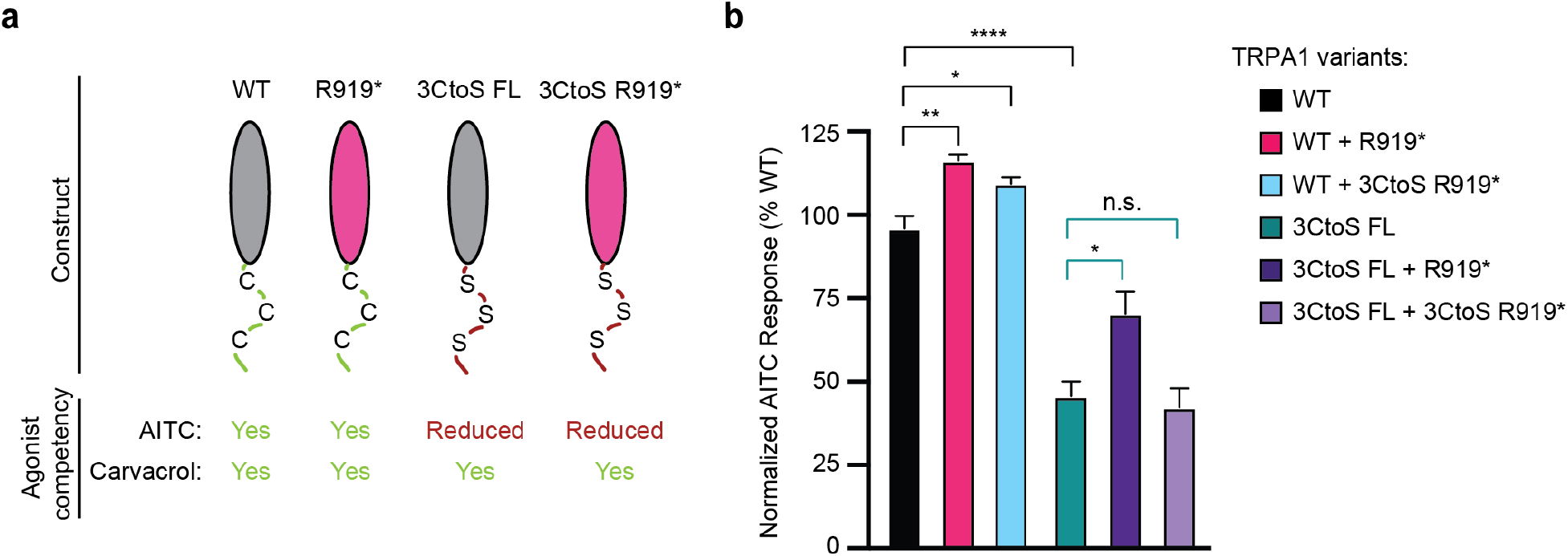
R919* TRPA1 subunits directly contribute to channel activity at the plasma membrane. (a) Schematic of TRPA1 constructs used for electrophile agonist complementation assays. Full-length (FL) hTRPA1 constructs indicated in grey, R919* hTRPA1 constructs indicated in pink. Agonist competency indicated in green (competent) or red (reduced competency). (b) Quantification of 10 μM AITC-evoked change in Fura-2 ratio relative to maximum response of each expression condition at 100 μM Carvacrol. Data further normalized to WT hTRPA1 as 100%. Data represent mean ± SEM. ****p<0.0001, **p<0.01, *p<0.05, n.s. not significant. n = 4 independent experiments, n ≥ 90 cells per transfection condition per experiment, one-way ANOVA with Bonferroni’s *post hoc* analysis.

### Source of R919* TRPA1-Conferred Hyperactivity

While AITC sensitivity of 3CtoS FL TRPA1 could only be restored by R919* TRPA1 subunits with intact cysteines, co-expression of WT TRPA1 with the R919* or 3CtoS R919* mutants conferred similar hyperactivity (Fig. 5b, compare black, pink, and blue bars). These results indicate the R919* mutant does not require intrinsic agonist sensitivity to modulate WT TRPA1 function and suggest that structural and/or regulatory features lost due to truncation contribute to channel sensitization.

TRPA1 structural features lost due to truncation in the R919* mutant can be subdivided into three discrete regions: the second pore helix and S6 transmembrane helix, the latter of which forms the lower gate; the TRP-like domain and subsequent ß-strand in the membrane-proximal cytoplasmic C-terminus as part of the allosteric nexus that play an active role in channel gating; and the remaining cytoplasmic C-terminus including an interfacial helix possibly involved in lipid regulation and the distal coiled coil, which forms the core intracellular inter-subunit interaction and coordinates a requisite polyanion cofactor (Fig. 1a-c)^15,16^. To analyze the contribution of each truncated region to the conferred channel sensitization, a suite of C-terminal truncation mutants was generated that exclude the coiled coil and interfacial helix (Δ1011), the complete cytoplasmic C-terminus (Δ969), and the S6 transmembrane helix and cytoplasmic C-terminus (Δ934) (Fig. 6a, Extended Data Fig. 9a and b). Akin to R919* TRPA1, these C-terminal truncations failed to form functional channels when expressed alone in HEK293T cells, suggesting that key regions in the distal cytoplasmic C-terminus are required to support proper channel function (Extended Data Fig. 9c). This inactivity is seemingly not due to poor expression or failure of the C-terminal truncations to traffic to the plasma membrane (Extended Data Fig. 9d). Each C-terminal truncation could immunoprecipitate WT TRPA1 when co-expressed, raising the possibility that they can also co-assemble with WT subunits (Fig. 6b). Ratiometric calcium imaging analysis of WT TRPA1 coexpressed with the partial (Δ1011) and complete (Δ969) cytoplasmic C-terminus truncations revealed a non-significant, but stepwise increase in AITC sensitivity (Fig. 6c, light and medium pink bars, Extended Data Fig. 10a). Additional truncation of the S6 transmembrane helix (Δ934) conferred robust and significant channel hyperactivity when co-expressed with WT TRPA1, exceeding that observed with the R919* mutant (Fig. 6c, dark pink bars, Extended Data Fig. 10a). Co-expression with the C-terminal truncation constructs affected WT TRPA1 expression in these experiments and thus AITC sensitivity was further normalized to WT TRPA1 expression (Extended Data Fig. 10b). These results suggest that truncation of the cytoplasmic C-terminus and the S6 transmembrane helix from TRPA1 subunits provides stepwise contributions to R919* TRPA1-conferred channel hyperactivity while additional removal of the second pore helix may weaken the conferred hyperactivity.

**Figure 6.**
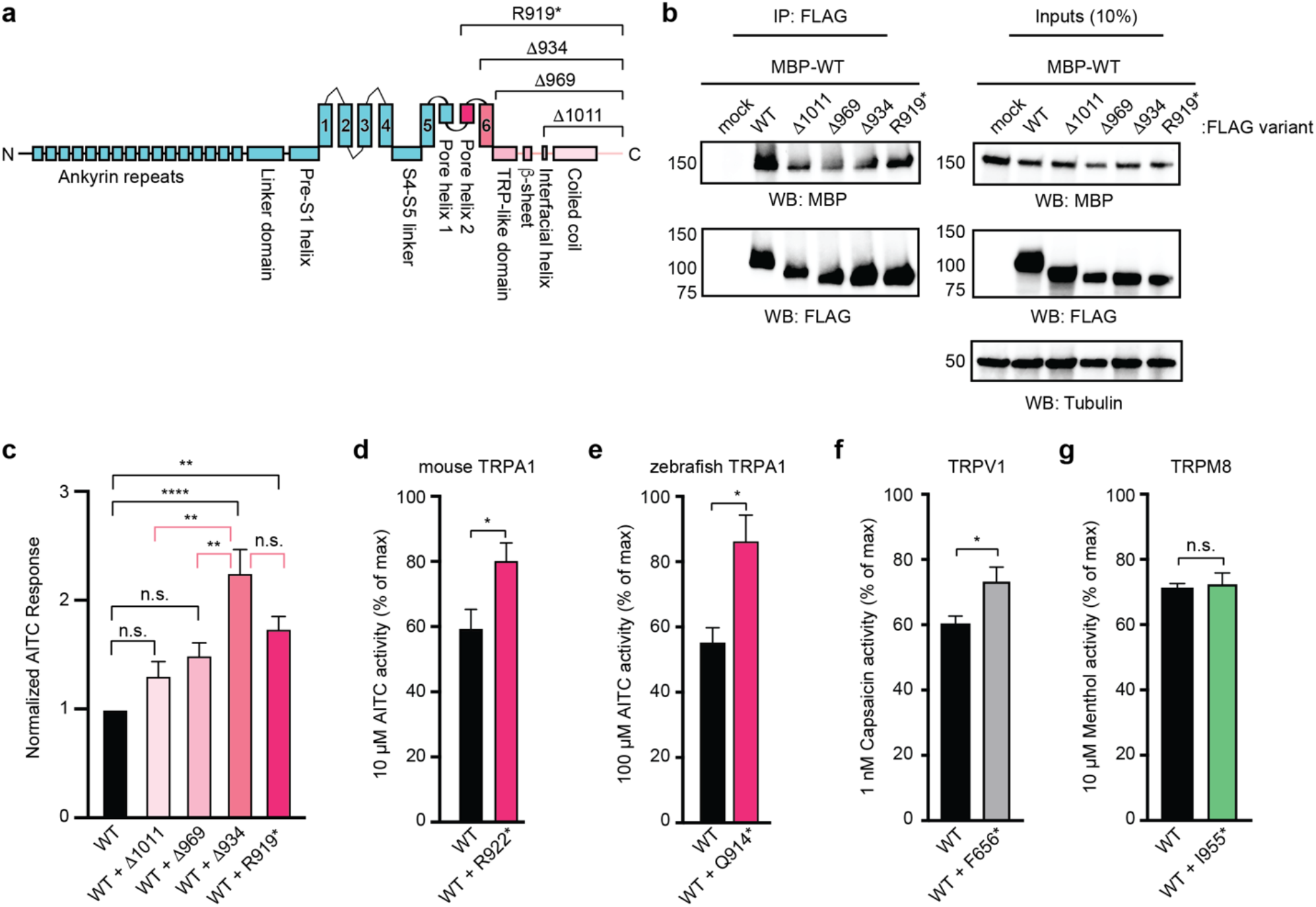
Characterization of the source, evolutionary conservation, and broad applicability of CFS-associated TRPA1 mutant-conferred channel hyperactivity. (a) Linear diagram depicting major structural domains in a WT hTRPA1 monomer. Missing portions from all C-terminal truncation constructs denoted above. R919* truncation completeness scales with gradations of pink. (b) Immunoblotting analysis of MBP-WT hTRPA1 protein expression after FLAG immunoprecipitation from lysates of HEK293T cells transiently co-transfected with empty vector (mock), or 3xFLAG-WT hTRPA1, 3xFLAG-R919* hTRPA1, or indicated 3xFLAG-hTRPA1 C-terminal truncation constructs. Samples were probed using anti-MBP antibody. 3xFLAG-tagged constructs were probed using HRP-conjugated anti-FLAG antibody. Tubulin from whole cell lysates (10%, inputs) was the loading control. Data is representative of 4 independent experiments. (c) Quantification of 10 μM AITC-evoked change in Fura-2 ratio relative to maximum response of each expression condition at 100 μM AITC. Data further normalized to WT hTRPA1 response and WT hTRPA1 expression. Colors as indicated in (A). Data represent mean ± SEM. **** p<0.0001, ** p<0.01, n.s. not significant. n = 4 independent experiments, n ≥ 60 cells per transfection condition per experiment, oneway ANOVA with Tukey’s *post hoc* analysis. (d) Quantification of 10 μM AITC-evoked change in Fura-2 ratio relative to maximum response of each expression condition with mouse TRPA1 variants to 100 μM AITC. Data represent mean ± SEM. *p<0.05. n = 4 independent experiments, n ≥ 90 cells per transfection condition per experiment, Student’s t-test. (e) Quantification of 100 μM AITC-evoked change in Fura-2 ratio relative to maximum response of each expression condition with zebrafish TRPA1 variants to 1 mM AITC. Data represent mean ± SEM. *p<0.05. n = 4 independent experiments, n ≥ 90 cells per transfection condition per experiment, Student’s t-test. (f) Quantification of 1 nM Capsaicin-evoked change in Fura-2 ratio relative to maximum response of each expression condition with human TRPV1 variants to 1 μM capsaicin. Data represent mean ± SEM. *p<0.05. n = 3 independent experiments, n ≥ 90 cells per transfection condition per experiment, Student’s t-test. (g) Quantification of 10 μM Menthol-evoked change in Fura-2 ratio relative to maximum response of each expression condition with rat TRPM8 variants to 200 μM Menthol. Data represent mean ± SEM. n.s. not significant. n = 3 independent experiments, n ≥ 90 cells per transfection condition per experiment, Student’s t-test.

### Conservation of CFS-Associated Mutant Conferred Hyperactivity and Broad Applicability

Negative stain electron microscopy of thirteen TRPA1 species orthologues has previously revealed a conserved channel architecture, which raises the possibility that the CFS-associated TRPA1 mutant may similarly confer channel hyperactivity in other orthologues^15^. The CFS-associated mutant was generated for mouse TRPA1 (R922*) and zebrafish TRPA1a (Q914*), which share 80% and 49% sequence identity to the human TRPA1 orthologue, respectively (Extended Data Fig. 11a). Expression of the mouse R922* or zebrafish Q914* TRPA1 mutants alone in HEK293T cells failed to form functional channels. However, they conferred significantly enhanced AITC sensitivity when co-expressed with their WT TRPA1 partners (Fig. 6d and e, Extended Data Fig. 11b and c). The ability of the mouse and zebrafish CFS-associated mutants to confer channel hyperactivity to the corresponding WT TRPA1 protein suggests that the evolutionary conservation of key structural features in the S6 transmembrane helix and cytoplasmic C-terminus serve to regulate channel activity.

Finally, the ability to selectively introduce TRPA1 hyperactivity in a genetically tractable manner with the CFS-associated mutant may represent a powerful and general approach to build similar genetic tools for other receptors. To test the feasibility, truncation mutants lacking the S6 transmembrane helix and the cytoplasmic C-terminus were generated for TRPV1 and the menthol receptor, TRPM8. Though members of the large TRP channel family, TRPA1, TRPV1 and TRPM8 have structurally distinct cytoplasmic domains, which make them great candidates for testing the broad applicability of this approach (Extended Data Fig. 12a). R919*-like mutants were not generated since TRPV1 and TRPM8 have elaborate extracellular pore domains that made it difficult to identify a comparable termination point. Like the analogous Δ936 TRPA1 truncation, neither F656* TRPV1 nor I955* TRPM8 formed functional channels in HEK293T cells despite expressing as well as their WT counterparts (Extended Data Fig. 12b-e). When co-expressed with their WT partners, F656* TRPV1 significantly enhanced capsaicin sensitivity (Fig. 6f and Extended Data Fig. 12b) while the I955* TRPM8 variant had no impact on menthol-evoked activity (Fig. 6g and Extended Data Fig. 12d). Thus, removal of the S6 transmembrane helix and cytoplasmic C-terminus similarly promotes TRPV1 channel sensitization, which is consistent with the enhanced channel activity conferred by an alternative splice variant that removes the part of the cytoplasmic C-terminus that mediates channel regulation by phosphatidylinositols^65,69^. In contrast, removal of these domains failed to confer channel hyperactivity to TRPM8, which is consistent with previous work that demonstrated components of the cytoplasmic C-terminus including the coiled coil are necessary for subunit oligomerization^49,70^. Collectively, these results illustrate analogous R919* or Δ936 TRPA1 mutants can be applied to other receptors to confer channel sensitization if their cytoplasmic C-terminal domains are not critical to proper channel assembly.

## DISCUSSION

Gain-of-function channelopathic mutations provide a unique opportunity to understand how changes in channel structure result in aberrant function. Most commonly, gain-of-function channelopathies introduce missense mutations in key structural regions that alter channel gating^23,27,28,71^. Far rarer are gain-of-function premature truncation channelopathies, of which only two have been previously characterized^48,72^. These characterized mutations truncate portions of the affected channel’s cytoplasmic C-terminal domain while retaining the full complement of transmembrane helices and the resulting proteins either exhibit aberrant channel activity or modulate the function of other receptors. In contrast, the CFS-associated R919* *TRPA1* mutation is a drastic nonsense channelopathic mutation, lacking core parts of the ion conduction pathway machinery in addition to the complete cytoplasmic C-terminus. Astonishingly, we found the R919* mutant conferred gain-of-function by enhancing agonist sensitivity when co-expressed with WT TRPA1 subunits. Our data support a model where WT and R919* TRPA1 subunits co-assemble to form hyperactive heteromeric ion channels, which represents a novel mechanism of gain-of-function by a truncation mutant (Fig. 7a). Our results indicate the R919* mutant sensitizes heteromeric channels due to loss of critical structural domains. Importantly, loss of these domains from all channel subunits *(e.g.,* in R919* TRPA1 homomers) yields nonfunctional complexes, revealing full channel architecture in at least one subunit is required to support channel functionality. Unexpectedly, we also find the R919* mutant can initiate channel activation with an electrophile agonist that targets the cytoplasmic domain despite lacking key machinery in the allosteric nexus. The R919* mutant, therefore, sheds light on the complex process of TRPA1 channel activation and provided us a unique opportunity to explore molecular mechanisms that govern channel activity.

**Figure 7.**
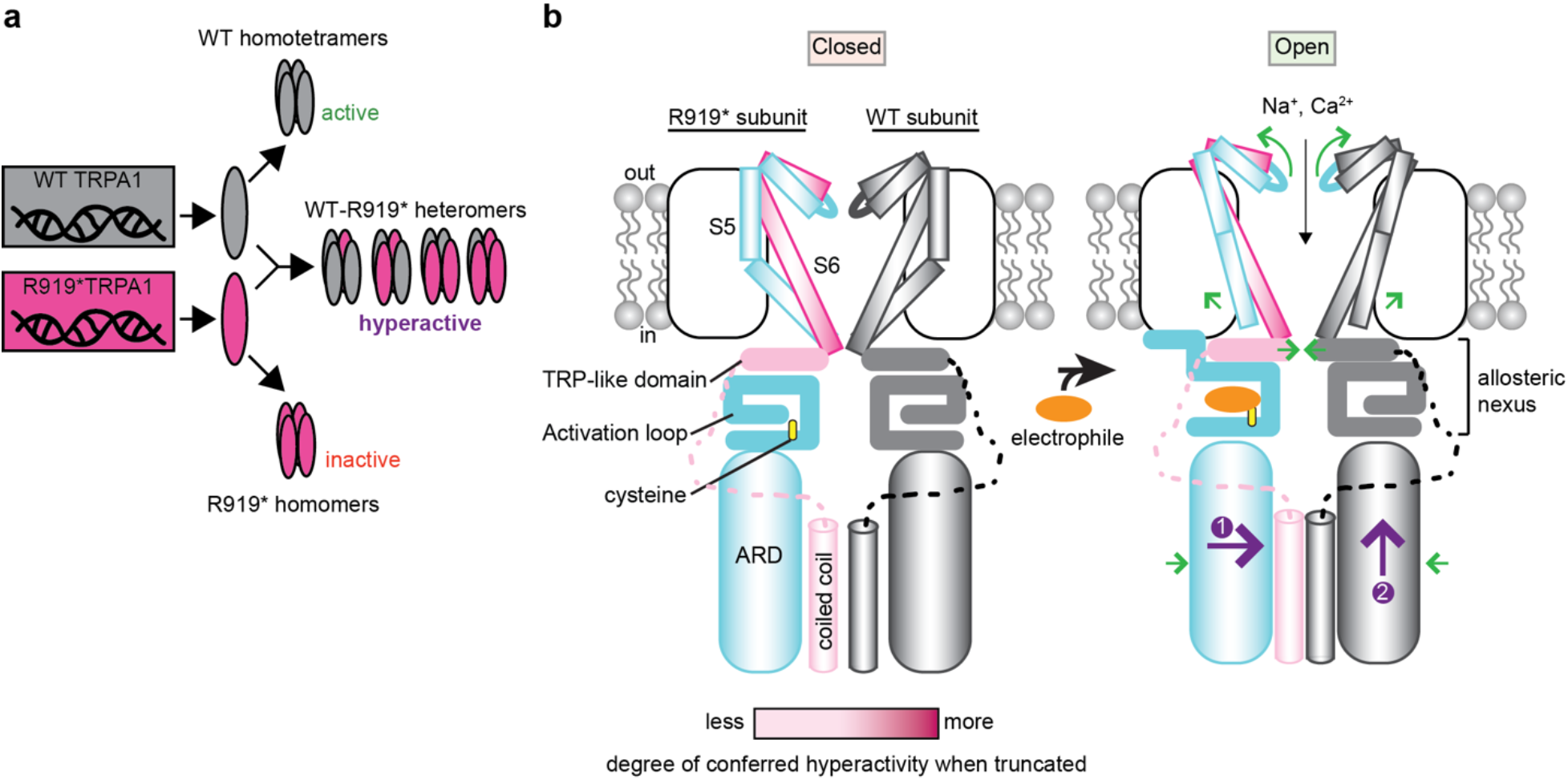
Mechanistic models of R919*-conferred channel hyperactivity. (a) Model of proposed mechanism of R919*-mediated TRPA1 sensitization. R919* TRPA1 patients are heterozygotes and carry WT and R919* *TRPA1* copies. WT and R919* hTRPA1 protein subunits can assemble into separate homomeric complexes that are active and inactive, respectively. WT and R919* hTRPA1 subunits may also co-assemble into hyperactive hetero-tetrameric channels of four possible subunit stoichiometries and configurations. We propose these heteromeric channels confer gain-of-function. (b) Model for R919* mutant-conferred channel sensitization and activation. In WT TRPA1 channels, electrophile agonist-evoked gating involves rearrangements in the cytoplasmic domains including conformational flipping of the activation loop, contraction of the coiled coil and adjacent ankyrin repeat domain (ARD), and a sliding rotation of the TRP-like domain towards the central channel axis (green arrows). These conformational changes occur with concerted dilation of the upper and lower gates in the S6 transmembrane helix and selectivity filter, which are coupled through straightening of the S5 transmembrane helix (green arrows). Loss of the cytoplasmic C-terminus and S6 transmembrane helix in the R919* mutant contribute to conferred hyperactivity in WT-R919* heteromer channels to different degrees (indicated by gradations of pink). Electrophile modification of reactive cysteines in R919* subunits (orange) may communicate channel activation to the pore through WT subunits *via* contraction of the ARD and coiled coil domains (step 1) that could propagate up to the allosteric nexus (step 2) (purple arrows). Two subunits are shown for clarity. Cartoon adapted from Zhao *et al*^17^ and Suo *et al*^16^.

High-resolution TRPA1 structures have been solved in distinct functional states, which allows us to hypothesize how loss of key structural features in the R919* truncation mutant may enhance channel activity^15–17^. Upon channel activation by an electrophile agonist, conformational changes originating in the cytoplasmic domains propagate to the transmembrane domain at the S5 and S6 helices (Fig. 7b, green arrows). Straightening of the S5 helix from one subunit initiates conformational changes in the S1-S4 voltage-sensor like domain of an adjacent subunit, promoting concerted channel activation through the domain-swapped architecture in the transmembrane domain^17^. While each subdomain lost in the R919* mutant undergoes gating-associated conformational changes that promote channel activation, these rearrangements may also serve as energetic barriers to gating when the channel is in the closed state. Indeed, C-terminal regions have been found to oppose gating in some ion channels^48,73–75^. Consistently, we found that loss of the cytoplasmic C-terminus conferred enhanced channel activity with a significant boost observed with additional loss of the S6 transmembrane helix (Fig. 7b, shades of pink). The R919* mutant subunit(s) did not need to be electrophile agonist competent to confer channel sensitization *(e.g.,* the R919* and 3CtoS R919* mutants conferred similar levels of sensitization, Fig. 5b). Therefore, we propose that loss of these domains in the R919* mutant provide release to the structural tension holding the S5 helix and the S1-S4 voltage sensor-like domain in specific conformations. This, in turn, may provide a stepwise reduction in the energetic barrier to channel activation in WT-R919* TRPA1 complexes. Interestingly, the ability of CFS-like mutants of two TRPA1 species orthologues to similarly confer gain-of-function indicates the structural and functional roles of the truncated domains are evolutionarily conserved.

Our agonist complementation assays revealed that the R919* mutant can initiate channel activation with electrophile agonists. This was surprising since the R919* subunits lack key structural features (*e.g.*, the TRP-like domain and the S6 helix) needed to communicate the electrophile agonist activation signal to the channel pore and to other subunits through the transmembrane domain (Fig. 7b). Thus, the R919* mutant subunits may be communicating the signal through an alternative mechanism. Upon channel activation, the membrane proximal ARD, which forms a cage around the coiled coil, contracts (Extended Data Fig. 13). Though R919* subunits lack the coiled coil helix, coiled coils can form with two or more helices and contraction of the R919* ARD may be “felt” by the coiled coil(s) of associated WT subunits (Fig. 7b, purple arrow 1). The coiled coil also contracts during channel activation, which may communicate the signal to the membrane proximal ARD and overlaying allosteric nexus of WT subunits (Fig. 7b, purple arrow 2). Indeed, coiled coil dynamics have been shown to play a critical role in regulating bacterial voltage-gated sodium channels and TRPM8^73,76^. Therefore, the membrane proximal ARD and coiled coil may represent an intracellular point of intersubunit communication that facilitates concerted channel gating akin to the transmembrane domain. This mechanism, in which ARD dynamics can contribute to channel gating, has exciting implications for how temperature sensing may be mediated by some TRPA1 species orthologues where the ARD has been shown to play a critical role in conferring or tuning thermosensation^20,21^. Structures of WT-R919* TRPA1 complexes are needed to directly reveal how R919* subunits alter channel architecture, which may provide a mechanism for channel activation through the mutant subunits.

The observation that R919* mutant subunits contribute to functional channels raises questions about how many R919* subunits are necessary to produce hyperactive complexes. Our tandem pulldown experiments crudely estimate the WT:R919* subunit composition of isolated heteromeric TRPA1 complexes as 1:1.3, which is consistent with formation of a mixture of WT:R919* subunit stoichiometries. This also suggests that if 2:2 WT:R919* assemblies do form, then statistically, these should be among the most abundant complexes. Future work is needed to determine which stoichiometries form in WT-R919* TRPA1 heteromers. Moreover, it is likely that only specific arrangements of WT-R919* TRPA1 complexes yield functional channels. Importantly, our activity assays were conducted by ratiometric calcium imaging, which reveals population-level channel function. It is possible that specific WT-R919* TRPA1 heteromer complexes have greatly altered channel properties that are masked or minimized in such ensemble experiments. Future electrophysiological work with a homogeneous channel population is needed to characterize specific WT-R919* TRPA1 heteromeric assemblies to determine which complexes are functional and to reveal altered channel properties such as ion selectivity, channel conductance, and activation or desensitization kinetics that may contribute to our observed channel hyperactivity.

Channel sensitization by the R919* mutant would be expected to prime TRPA1 channels for activation and may lower the barrier to initiate pain signals and neurogenic inflammation. Thus, this gain-of-function could account for the clinical phenotype of neuronal hyperexcitability-hypersensitivity observed in CFS patients harboring the R919* *TRPA1* mutation. The R919* mutant represents a genetically tractable method to selectively confer TRPA1 sensitization, which will facilitate future work in mouse models and isolated neurons to determine the behavioral and physiological consequences of chronic TRPA1 hyperactivity. Such models may additionally be helpful to identifying and vetting analgesic and anti-inflammatory agents to treat pain and other pathological conditions associated with aberrant TRPA1 activity.

Our discovery that the R919* TRPA1 mutant confers gain-of-function when co-expressed with WT subunits is noteworthy and distinct from the dominant negative or loss-of-function phenotypes observed with most other characterized nonsense channelopathic mutations^45,77–79^. Indeed, a *TRPA1* knockout allele that truncates the resulting protein after the S4 transmembrane helix ablates TRPA1 activity in homozygous mice and confers haploinsufficiency in heterozygous animals^26^. Thus, it is remarkable that the R919* TRPA1 mutant, which contains one additional transmembrane helix than the *TRPA1* knockout allele, confers gain-of-function. Our findings expand the possible physiological impact of seemingly nonfunctional truncation mutants, alternative splice variants, or other single nucleotide polymorphisms that are endogenously coexpressed with WT protein and demonstrate that it is critical to consider the genotype of patients where such gene permutations are identified. Finally, our ability to recapitulate channel sensitization with a TRPV1 truncation mutant raises the possibility that such mutations may serve as a novel and general approach to similarly probe channel structure and function with experimental and naturally occurring deletions.

## EXPERIMENTAL PROCEDURES

### Plasmid Construction

DNA sequences for human TRPA1, human TRPV1, and rat TRPM8 were amplified by PCR and subcloned into p3xFLAG-eYFP-CMV-7.1 vector (Addgene) at the NotI/BamHI sites or into 8xHis-MBP pFastBac1 modified with a CMV promoter (obtained from David Julius’ lab) at the BamHI/NotI sites using In-Fusion EcoDry cloning (Takara) according to manufacturer protocols. Mouse TRPA1 and zebrafish TRPA1a orthologs were similarly subcloned into p3xFLAG-eYFP-CMV-7.1. For most calcium imaging experiments, untagged human TRPA1 in a combined mammalian/oocyte expression vector pMO (modified from pcDNA3 - obtained from David Julius’ lab) was used. For the calcium imaging experiments presented in Fig.s 6C-E and Extended Data Fig. 3C, 8C, and 9, 3xFLAG-tagged TRPA1 constructs were used. WT and inactive human TRPA1 variants were subcloned into CaM/pIRES2-eGFP (Addgene) at the NheI/EcoRI sites to generate a positive fluorescent readout for transfection in singly transfected calcium imaging studies. BioID2-tagged variants were generated by introducing a BamHI site between the 8xHis and MBP tags in the 8xHis-MBP-pFastBac1 modified with a CMV promoter vector by Quikchange Lightning site-directed mutagenesis (Agilent) and cloning BioID2 into the BamHI sites using In-Fusion EcoDry cloning according to manufacturer protocols.

Point mutations to generate the CFS-associated mutant premature stop codon (R919* for human TRPA1, R922* for mouse TRPA1, and Q914* for zebrafish TRPA1a), truncation constructs for TRPV1 (F656*) and TRPM8 (I955*), and electrophile incompetent variants were implemented using Quikchange Lightning site-directed mutagenesis. Larger deletions of C-terminal domains of TRPA1 were accomplished via InFusion EcoDry cloning. For tandem purification studies, a Strep-tag II (Trp-Ser-His-Pro-Gln-Phe-Glu-Lys) or a FLAG tag (Asp-Tyr-Lys-Asp-Asp-Asp-Asp-Lys) was added on the N-terminus of MBP WT or R919* TRPA1 using Quikchange Lightning site-directed mutagenesis (Agilent).

All DNA primers were ordered from ThermoFisher and all constructs were sequence-verified using the Yale School of Medicine Keck DNA Sequencing Facility.

### Cell Culture and Protein Expression

Human embryonic kidney cells (HEK293T, ATCC CRL-3216) were cultured in Dulbecco’s modified Eagle’s medium (DMEM; Invitrogen) supplemented with 10% calf serum for all experiments except immunofluorescence imaging where 10% FBS was used, and 1x Penicillin-Streptomycin (Invitrogen) at 37°C and 5% CO2. Cells were grown to ~85-95% confluence before splitting for experiments or propagation. HEK293T cells cultured to ~95% confluence were seeded at 1:10 or 1:20 into 6- or 12-well plates (Corning), respectively. After 1-5 hours recovery, cells were transiently transfected with 1 or 0.5 μg plasmid for 6- or 12-well plates, respectively using jetPRIME (Polyplus) according to manufacturer protocols. Total transfected DNA per well was normalized with empty pMO vector, as necessary.

### Ratiometric Calcium Imaging

40-48 hours post-transfection, HEK293T cells were plated into isolated silicone wells on poly-L-lysine-coated cover glass. 1 hour later, cells were loaded with 10 μg/mL Fura 2-AM (Sigma) in Ringer’s solution (Boston Bioproducts) and incubated for 45 min at 37°C, then rinsed twice with Ringer’s solution. Ratiometric calcium imaging was performed using a Zeiss Axio Observer 7 inverted microscope with a Hamamatsu Flash sCMOS camera at 20x objective. Dual images (340 and 380 nm excitation, 510 nm emission) were collected and pseudocolour ratiometric images were monitored during the experiment (Metafluor software). After stimulation with agonist, cells were observed for 45-100 s. AITC, Carvacrol, Capsaicin, Menthol, and Cinnamaldehyde were all purchased from Sigma and were freshly prepared as stocks at 4x the desired concentration in 1% DMSO and Ringer’s solution. 5 μL 4x agonist was added to wells containing 15 μL Ringer’s solution to give the final 1x desired concentration. For the antagonist experiments in Fig. 2C and Extended Data Fig. 2, cells were pre-treated with Ringer’s solution containing A-967079 (Tocris), HC-030031 (Tocris), or Ruthenium red (Sigma) at the desired concentration for 1 min and activated with a 4x AITC solution containing 4x antagonist to maintain antagonist concentration. When quantifying calcium influx responses for dose-response curves, each experimental replicate at each agonist concentration was an average response from 30 cells, with three separate replicates *(e.g.,* 90 total cells per agonist concentration). For all other experiments, a minimum of 60-90 cells were selected per condition per replicate for ratiometric fluorescence quantification in Metafluor with 3-5 replicates per experiment. Preactive cells were excluded from quantification and background signal was quantified from un-transfected cells and subtracted from quantified cells for normalization. Unless reported in arbitrary units (AU), all data were normalized by the maximum response evoked for that transfection condition by 10x agonist (100 uM for human and mouse TRPA1, and 1 mM for zebrafish TRPA1), 200x agonist (200 μM Menthol), or 1000x agonist (1 μM Capsaicin). For experiments presented in Fig. 6C-E and Extended Data Fig. 4C, 9C, 10, and 12, the imaged cells were lysed, and quantitatively immunoblotted for 3xFLAG-tagged TRPA1 to ensure equivalent expression, as detailed below. For inactive variants or antagonist-treated TRPA1 that did not exhibit appreciable calcium influx, constructs in the pIRES-eGFP vector were transfected in HEK293T cells. Fluorescence images were collected with a GFP filter after ratiometric calcium imaging, which allowed selection of GFP positive cells for subsequent quantification. R919* hTRPA1 data in Fig. 1E was quantified as above. Data in Fig. 2C represent the percentage inhibition of the AITC-evoked maximal calcium influx for mock-treated WT hTRPA1.

### Immunofluorescence Imaging

HEK293T cells were transiently transfected with hTRPA1 constructs using Lipofectamine 2000 transfection reagent (ThermoFisher) according to manufacturer’s protocol and incubated for 20 hrs prior to immunostaining. Cells were fixed on coverslips with 3.5% paraformaldehyde for 15 minutes, and then incubated with 5 ug/mL Alexa Fluor 350-conjugated Wheat Germ Agglutinin (Invitrogen) in PBS at room temperature for 10 minutes, followed by permeabilization using PBS + 0.5% Triton X-100 for 10 minutes and incubation with PBS + 1% Bovine serum albumin (BSA) for 10 minutes. Cells were incubated with primary antibodies diluted 1:1000 in PBS + 1% BSA for 16 hours at 4°C. Cells were washed with PBS and incubated with secondary antibodies diluted 1:1000 in PBS + 1% BSA for 1 hour at room temperature. Images were acquired on a Nikon ECLIPSE Ti2 with a Hamamatsu Fusion sCMOS camera at 100X magnification.

### Image Analysis and Deconvolution

For quantitative analysis of R919* TRPA1 intensity at the plasma membrane, raw single stack immunofluorescence images were processed in ImageJ using the line scanning function. A straight line was drawn through the width of at least 30 cells and the fluorescence intensity of each pixel was measured as a function of distance for each channel. Intensity was normalized to the average middle 30% of each line scan representing the cell interior, and distance was normalized from 0 to 1 to account for differences in cellular size. Line scans were compiled using GraphPad Prism, and the average maxima from all scans was taken to represent fluorescence intensity at the plasma membrane. For quantitative analysis of WT and R919* TRPA1 co-localization, raw single stack immunofluorescence images were processed in ImageJ using the line scanning function. Freehand lines were drawn over the plasma membrane, and the intensity of at least 450 pixels were measured for each channel. Measurements were normalized to the average intensity of each line scan to account for variability in transfection efficiency. Pearson’s correlation coefficient (r) and the coefficients of determination (r^2^) were calculated to represent WT and R919* TRPA1 colocalization. To deconvolve immunofluorescence images, stacks of optical sections (z step = 0.2 μm) were restored with Huygens software (Scientific Volume Imaging) using the maximum-likelihood estimation algorithm. The restored stacks were further processed in Photoshop CS4 (Adobe).

### Cell Lysis and Pulldown Experiments

40-48 hours post-transfection, HEK293T cells were lysed in 100-200 μL of <u>TRPA1 lysis buffer</u> (20 mM HEPES, 150 mM NaCl, 1 mM IP_6_, 20 mM DDM, 1x EDTA-free Roche Complete Protease Inhibitor Cocktail, pH 7.8) on ice for 10 min with light mechanical agitation and collected by gentle pipetting. Resulting lysates were rotated for an additional 10 minutes at 4°C, followed by centrifugation at 15,000 rpm for 10 min at 4°C. Total protein concentration in lysates was quantitated using BCA assay (Pierce). Equal concentrations of protein lysate from each experimental condition were added to affinity resins, as specified below. 10% of loaded protein amount was reserved as a whole-cell lysate loading control.

#### FLAG Immunoprecipitation

Lysates were incubated with 5-7.5 μL of lysis buffer-equilibrated EZview Red Anti-FLAG M2 affinity resin (Sigma) for 1 hour at 4°C with gentle rotation. Resin beads were washed four times with lysis buffer prior to elution; resin was pelleted, and associated proteins were eluted in 125 μg/ml 3xFLAG peptide (Sigma) for 30 minutes on ice.

#### Amylose Pulldown

Lysates were incubated with 15 μL of lysis buffer-equilibrated amylose resin (New England Biolabs) for 1 hour at 4°C with gentle rotation. The resin beads were washed five to six times with lysis buffer prior to elution; resin was pelleted, and MBP-tagged and associated proteins were eluted in 60 mM maltose (Sigma) for 30 minutes on ice.

#### Strep Pulldown

Lysates were incubated with 12.5 μL of lysis buffer equilibrated Strep Tactin Sepharose (IBA) for 1 hour at 4°C with gentle rotation. The resin beads were washed six times with lysis buffer prior to elution; resin was pelleted, and strep-tagged and associated proteins were eluted in 1X Buffer E (IBA) diluted in lysis buffer for 30 minutes on ice. For tandem pulldown assays, strep-enriched eluates were subsequently incubated with 10 μL of lysis buffer-equilibrated EZview Red Anti-FLAG M2 affinity resin for 2 hours at 4°C with gentle rotation and washed 6 times with lysis buffer prior to elution as above.

#### Neutravidin Pulldown

Cell lysates that were generated following surface labeling or proximity biotinylation experiments were incubated with 15 μL of lysis buffer equilibrated Neutravidin resin (Pierce) for 2 hours at 4°C with gentle rotation. The resin was then washed with lysis buffer three times, followed by a harsher wash with 1x PBS + 100 mM DTT. Resin was then washed once each with lysis buffer and 1x PBS. Biotinylated protein was eluted from the resin with a multi-step protocol to prevent TRPA1 aggregation while maximizing protein elution from the resin. First, resin was incubated with 10 μL of <u>biotin elution buffer</u> (TRPA1 lysis buffer, 100 mM Glycine, 10 mM Biotin, 1% SDS) on ice for 10 min, followed by addition of 1 μL ß-mercaptoethanol (BME; Sigma) to each sample and incubation on ice for 5 minutes, and finally by addition of 4 μL 4x Laemmli buffer + 10% BME with incubation at 65°C for 10 minutes. The resin was centrifuged, supernatant was removed and combined with additional 4 μL Laemmli buffer + 10% BME for SDS-PAGE analysis.

### SDS-PAGE and Immunoblot

Samples were combined with Laemmli Sample buffer + 10% BME and separated on pre-cast 4-20% SDS-PAGE gels (Bio Rad). Gels were transferred onto PVDF membranes (BioRad) by semi-dry transferred at 15V for 40 minutes. Blots were blocked in 3% BSA prior to antibody probing. The following primary antibodies were used in PBST buffer (Boston Bioproducts): MBP (mouse, 1:30,000, New England Biolabs), FLAG (mouse, 1:30,000, Sigma), strep (mouse, 1:30,000, IBA), tubulin (mouse, 1:5,000 in BSA, Sigma). HRP-conjugated IgG secondary anti-mouse antibody was used as needed (rabbit, 1:25,000, Invitrogen). Membranes were developed using Clarity Western ECL substrate (Bio-Rad) and imaged using a Chemidoc Imaging System (BioRad). Densitometric quantifications were performed with ImageJ software. All quantified band intensities for eluted samples were divided by their tubulin-normalized input band intensities, except for the tandem purification crude stoichiometric analysis in Fig. 4F - where only MBP signal intensity in eluates was used.

### Surface Biotinylation Assay

Surface biotinylation assays were adapted from previously reported protocols^80^. Briefly, HEK293T cells were seeded in a 6-well plate pre-coated with Poly-L-Lysine (Sigma) and transfected with expression plasmids. 40-48 hours post-transfection, cells were washed with PBS, placed on ice, and incubated for 20 minutes with chilled 0.05 mg/mL Sulfo-NHS-LC-Biotin (ThermoFisher) in Ringer’s solution. Cells were then washed with chilled wash buffer (PBS + 100 mM Glycine) and the reaction was quenched on ice for 30 minutes with 100 mM Glycine and 0.5% BSA in PBS. Cells were then washed three times and lysed in TRPA1 lysis buffer containing 100 mM Glycine directly on the 6-well plate. Lysates were collected, protein concentration was determined by BCA assay, biotinylated proteins were isolated by Neutravidin pulldown and analyzed by immunoblot as described above. During quantification, any FLAG signal in the torsin eluates lane was subtracted from all other conditions to account for probe internalization.

### Proximity Biotinylation Assays

BioID2- or 3x-FLAG-tagged TRPA1 and Kv1.2/2.1 fusion proteins were co-expressed in HEK293T cells. ~24 hours post-transfection, the cell culture media was supplemented with 50 uM biotin. The following day, cells were lysed with TRPA1 lysis buffer and subjected to affinity purification using neutravidin pulldowns, followed by a immunoblot, as described above.

### Fluorescence Size Exclusion Chromatography (FSEC)

eGFP- or 3x-FLAG-tagged TRPA1 fusion proteins were co-expressed in HEK293T cells (3 μg total DNA/well); 1 well was transfected for the WT-only condition and 3 wells were transfected for R919*-containing conditions to account for lower expression. 40-48 hours post-transfection, 500 (WT/WT), 1,650 (WT/R919*), and 2,700 (R919*/R919*) μg of lysate were subjected to FLAG immunoprecipitation as described above. FSEC procedures were adapted from those previously reported^15^. Briefly, sample volumes were brought to 1 mL with TRPA1 lysis buffer, then passed through 0.22-micron filters (Costar) by centrifugation, loaded onto a Superose 6 Increase column (GE Healthcare) pre-equilibrated with <u>FSEC buffer</u> (20 mM HEPES, 150 mM NaCl, 1 mM BME, 0.5 mM DDM, 1 mM IP_6_, pH 8), and run at a flow rate of 0.5 ml/min. The in-line fluorescence detector (Shimadzu) settings were as follows: excitation, 488 nm; emission, 510 nm; time increment, 1 s; integration time, 1 s. Fractions corresponding to TRPA1 and free eGFP peaks were collected, concentrated using Amicon Ultra 10 KDa cut-off spin filters, and subjected to immunoblot as described above.

### Blue Native PAGE

Lysates containing dual-tagged WT and R919* TRPA1 were tandem-purified with strep and FLAG affinity resins as described above. Eluates were split into two aliquots, one of which was given full denaturation treatment in Laemmli Sample buffer + 10% BME, while the second was added to NativePAGE Sample Buffer and 5% G-250 Sample Additive (Invitrogen) as per manufacturer instructions. Samples were loaded onto 4–15% Mini-Protean TGX Gels (Bio-Rad) and run at 180 V in NativePAGE Running Buffer (Invitrogen) with 10x Coomassie additive in the cathode buffer. After 25 min, the voltage was adjusted to 115 V for the remainder of the run. After 40 min of run time, the 10x Coomassie cathode buffer was removed and replaced with 1x Coomassie running buffer for the remaining 1.5 hours. Transfer to PVDF membrane and immunoblotting were performed as described above.

### Statistical Analysis

All data quantification was performed in Microsoft Excel. Quantified data presentation and statistical analyses were performed in GraphPad Prism. The GraphPad Prism colorblind safe color pallet was applied to all quantified data presented. Criterion for statistical significance for all tests was p<0.05.

## ACKNOWLEDGEMENTS

We thank Wendy Gilbert, Franziska Bleichert, Elena Gracheva, Joe Howard, Michael Koelle, Christian Schlieker, Tony Koleske, Yong Xiong, and Mark Solomon for constructive suggestions, Grover Paulsen-Sharpe for moral support, and members of the Paulsen lab for helpful discussions and for critical reading of the manuscript. We, additionally, gratefully acknowledge Christian Schlieker for providing the *FLAG-Torsin A* plasmid. A.B. is supported by a Biophysics pre-doctoral training grant (5T32GM008283-33). This research was supported by an International Association for the Study of Pain Early Career Research Grant, a Rita Allen Foundation and American Pain Society Pain Scholar Award, and by NIH grant R35GM142825 to C.E.P., by NIH grant CA244865 to L.K., and by the Pershing Square Sohn Cancer Research Alliance to I.T. and L.K. The content is solely the responsibility of the authors and does not necessarily represent the official views of the National Institutes of Health.

A.B., S.P.S., and C.E.P. planned the project. A.B., S.P.S., I.T., and C.E.P. designed experiments. A.B., S.P.S., A.L.Z., and C.E.P. carried out calcium imaging and surface biotinylation. A.B., S.P.S., and C.E.P. carried out pulldowns and proximity biotinylation. A.B. performed Blue Native PAGE. S.P.S. carried out fluorescence size exclusion chromatography and crude subunit stoichiometry. I.T. carried out immunostainings. A.B., S.P.S., and C.E.P. wrote the manuscript with input from I.T., A.L.Z., and L.K.

## Notes

### Competing Interest Statement

The authors have declared no competing interest.

