## Supplemental Material for "Molecular Mechanism of Hyperactivation Conferred by a Truncated TRPA1 Disease Mutant Suggests New Gating Insights"

### SUPPLEMENTARY INFORMATION

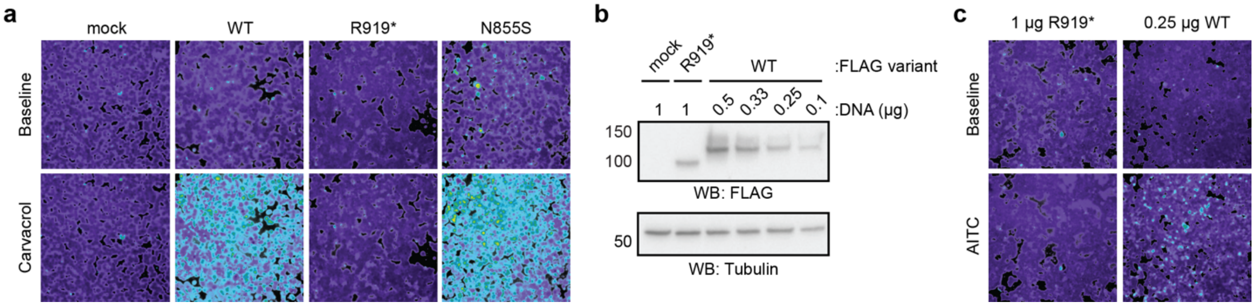


**Extended Data Figure 1. The R919* mutant is a nonfunctional TRPA1 natural variant.**
(a) Ratiometric calcium imaging of HEK293T cells transiently transfected with empty vector (mock), WT hTRPA1, R919* hTRPA1, or N855S hTRPA1. Cells were stimulated with Carvacrol (100 µM). Images are representative of three independent experiments. (b) Western blot of lysates from transiently transfected HEK293T cells expressing 3xFLAG-tagged hTRPA1 variants, probed as in (Fig. 1g). HEK293T cells were transfected with the indicated amount of plasmid (in µg). (c) Ratiometric calcium imaging of HEK293T cells transiently transfected with the indicated amount of 3xFLAG-tagged WT or R919* hTRPA1 to achieve comparable expression levels. Cells were stimulated with AITC (100 µM). Images are representative of three independent experiments.

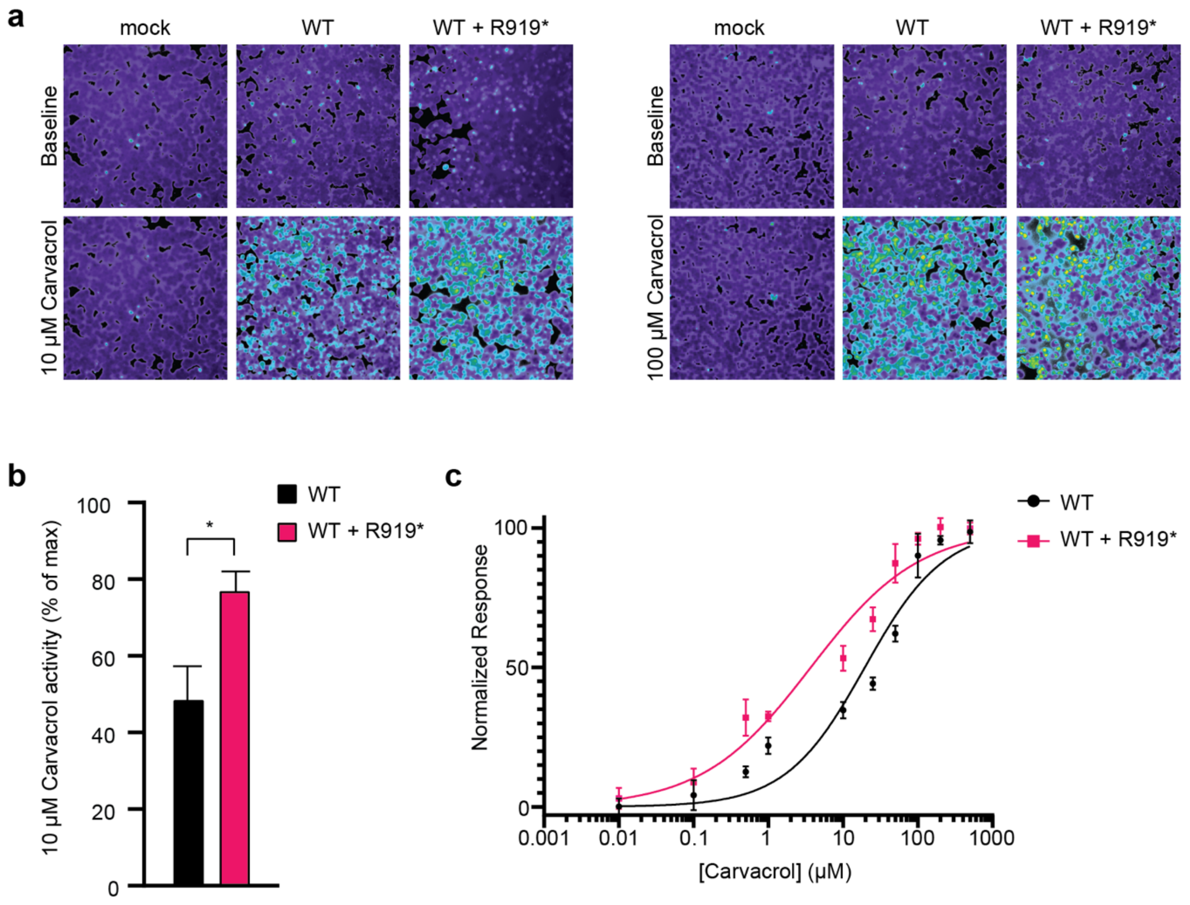


**Extended Data Figure 2. The R919* mutant confers hyperactivity when co-expressed with WT TRPA1 subunits.** (a) Ratiometric calcium imaging of HEK293T cells transiently transfected with empty vector (mock), WT hTRPA1, or WT and R919* hTRPA1. Cells were stimulated with Carvacrol (10 or 100 µM). Images are representatives from three independent experiments. (b) Quantification of 10 µM Carvacrol-evoked change in Fura-2 ratio relative to maximum response of each expression condition at 100 µM Carvacrol. Data represent mean ± SEM. *$p<0.05$. n = 4 independent experiments, n ≥ 90 cells per transfection condition per experiment, Student's t-test. (c) Dose-response curve of Carvacrol-evoked calcium responses for HEK293T cells transiently transfected with WT hTRPA1 or WT and R919* hTRPA1. Calcium responses normalized to maximum calcium response to 500 µM Carvacrol. Traces represent the average ± SEM of normalized calcium responses from 3 independent experiments, n = 30 cells per agonist concentration per experiment. Data were fit to a non-linear regression. $EC_{50}$ (95% CI) values are 19.5 µM for WT hTRPA1 (95% CI, 9.6-33.8 µM) and 3.7 µM for WT and R919* hTRPA1 (95% CI, 1.9-6.8 µM). Carvacrol $EC_{50}$ reduction for WT and R919* hTRPA1 co-expression is statistically significant ($p<0.001$, Extra sum-of-squares F test).

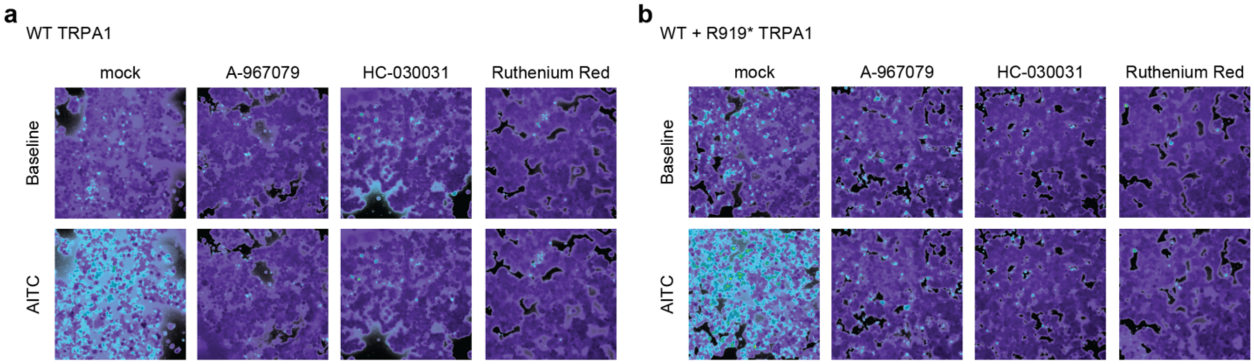


**Extended Data Figure 3. WT and WT + R919* channels are inhibited by canonical TRPA1 antagonists and pore blockers.** (a-b) Ratiometric calcium imaging of HEK293T cells transiently transfected with (a) WT hTRPA1, or (b) WT and R919* hTRPA1 from data quantified in Figure 2c. Cells were pre-treated with Ringer's solution (mock), A-967079 (10 μM), HC-030031 (30 μM), or Ruthenium Red (10 μM). Channels were activated with AITC (100 μM). Images are representatives from three replicate measurements.

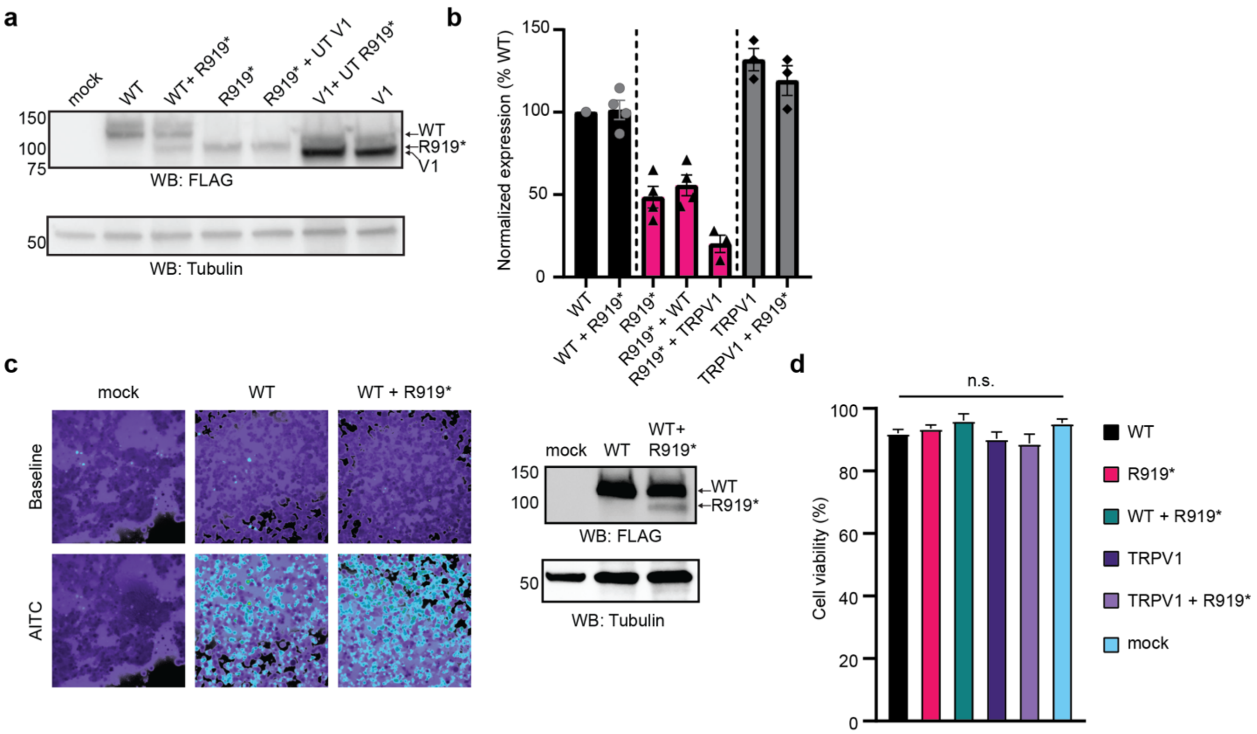


**Extended Data Figure 4. The R919* mutant does not affect WT hTRPA1 or TRPV1 expression levels and does not cause general cell stress.** (a) Western blot of lysates from transiently transfected HEK293T cells expressing empty vector (mock), 3xFLAG-WT hTRPA1, 3xFLAG-WT and R919* hTRPA1, 3xFLAG-R919* hTRPA1, 3xFLAG-R919* hTRPA1 and untagged (UT) hTRPV1, 3xFLAG-hTRPV1 and untagged (UT) R919* hTRPA1, or 3xFLAG-hTRPV1. Lysates were probed using HRP-conjugated anti-FLAG antibody. Tubulin was the loading control. (b) Quantitative analysis of Tubulin-normalized 3xFLAG-tagged hTRPA1 variants or hTRPV1 from (A) relative to WT hTRPA1. Data represent mean ± SEM. n=3-4 independent experiments. (c) Ratiometric calcium imaging (left) and Western blot analysis (right) of HEK293T cells transiently transfected with empty vector (mock), 3xFLAG-WT hTRPA1, or 3xFLAG-WT and R919* hTRPA1. Cells were stimulated with AITC (10 μM). Images are representatives from three replicate measurements. Lysates were probed using HRP-conjugated anti-FLAG antibody. Tubulin was the loading control. (d) Cell viability of HEK293T cells transiently transfected with the indicated hTRPA1 variants or hTRPV1 was quantified by trypan blue exclusion. Data represent mean ± SEM. n=4-17 independent readings per transfection condition, one-way ANOVA with Bonferroni's *post hoc* analysis.

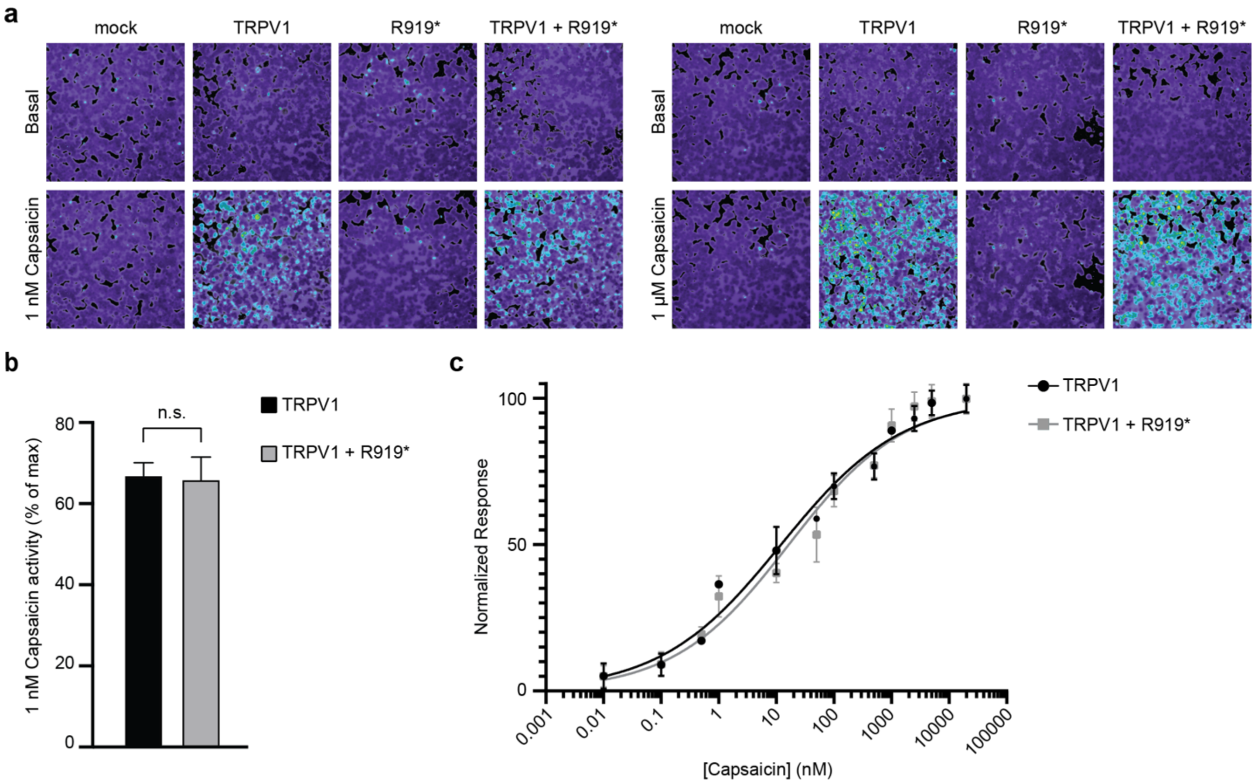


**Extended Data Figure 5. The R919* mutant does not affect TRPV1 channel activity.** (a) Ratiometric calcium imaging of HEK293T cells transiently transfected with empty vector (mock), hTRPV1, R919* hTRPA1, or hTRPV1 and R919* hTRPA1. Cells were stimulated with 1 nM Capsaicin (left) or 1 µM Capsaicin (right). Images are representatives from three independent experiments. (b) Quantification of 1 nM Capsaicin-evoked change in Fura-2 ratio relative to maximum response of each expression condition at 1 µM Capsaicin. Data represent mean ± SEM. n.s., not significant. n = 3 independent experiments, n ≥ 90 cells per transfection condition per experiment, Student's t-test. (c) Dose-response curve of Capsaicin-evoked calcium responses for HEK293T cells transiently transfected with hTRPV1 or hTRPV1 and R919* hTRPA1. Calcium responses normalized to maximum calcium response at 20 µM Capsaicin. Traces represent the average ± SEM of normalized calcium responses from 3 independent experiments, n = 30 cells per agonist concentration per experiment. Data were fit to a non-linear regression. $EC_{50}$ (95% CI) values are 12.1 nM for hTRPV1 (95% CI: 7.3-19.7 nM) and 16.5 nM for hTRPV1 and R919* hTRPA1 (95% CI: 9.3-28.6 nM).

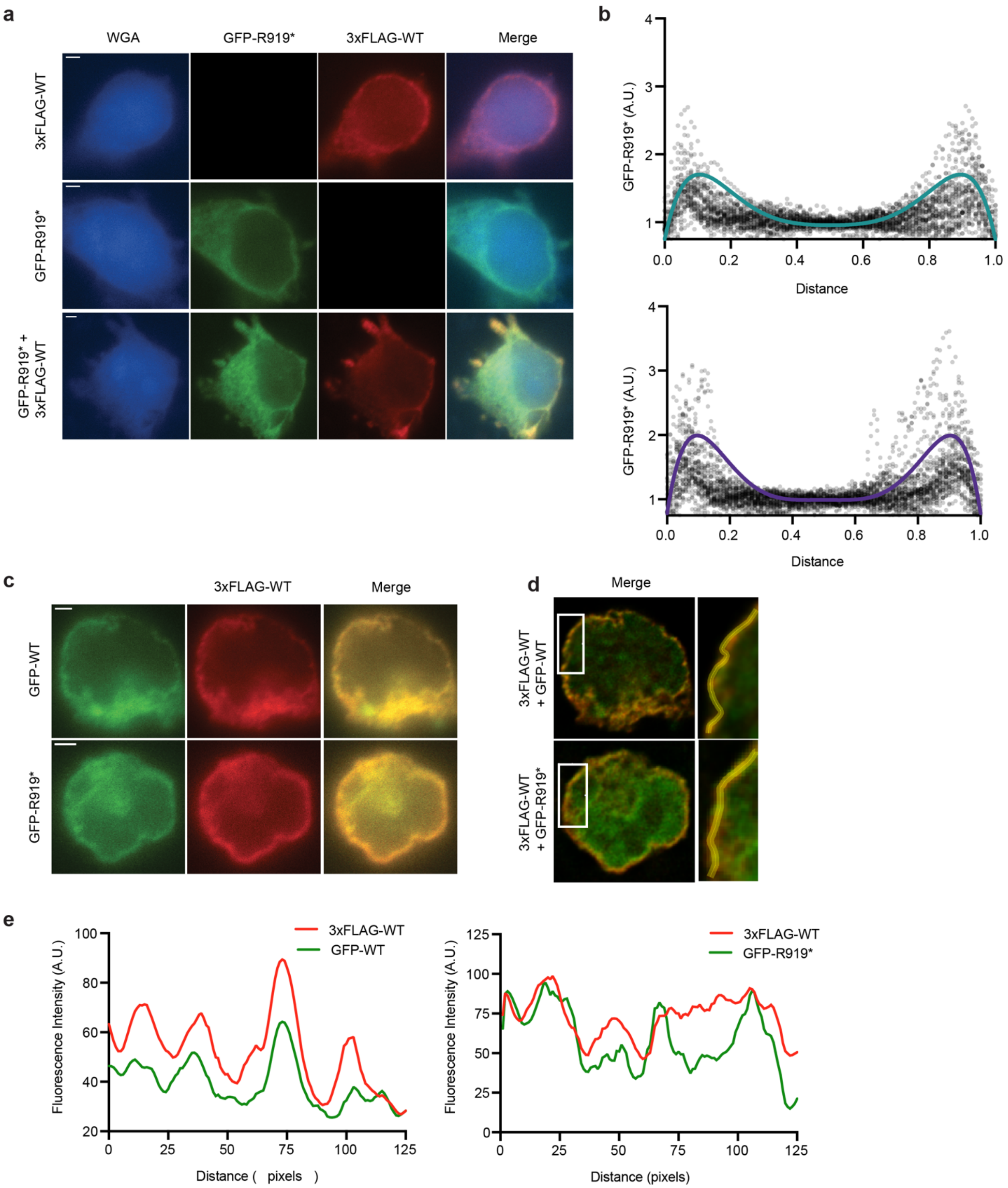


**Extended Data Figure 6. Representative immunofluorescence imaging data.**
(a) Representative raw immunofluorescence images of HEK293T cells transiently transfected with GFP-R919* hTRPA1, 3xFLAG-WT hTRPA1, or GFP-R919* hTRPA1 and 3xFLAG-WT hTRPA1. Cells were stained with anti-GFP (green) and anti-FLAG (red) antibodies. Plasma membrane was labeled with wheat germ agglutinin (blue). Scale bar indicates 2 µm. (b) Combined line scans of HEK293T cell cross-sections with transient transfection of GFP-R919* hTRPA1 (left) or co-transfection of GFP-R919* hTRPA1 and 3xFLAG-WT hTRPA1 (right). Distance and GFP-R919* hTRPA1 fluorescence are normalized relative to

total cell width and internal intensity. Representative polynomial lines are overlayed in teal (left) or purple (right). n=30 cells. (c) Raw images of transiently transfected HEK293T cells used for fluorescence correlation in Figure 3. Cells were transiently transfected with 3xFLAG-WT hTRPA1 and GFP-WT hTRPA1 or GFP-R919* hTRPA1, then stained with anti-GFP (green) and anti-FLAG (red) antibodies. Scale bar indicates 2 μm. (d) Deconvolved images of HEK293T cells depicted in (c). Segments of plasma membrane used are magnified (right) for line-scan analysis of red and green signal intensity. (e) Line scans of plasma membrane segments indicated in (d).

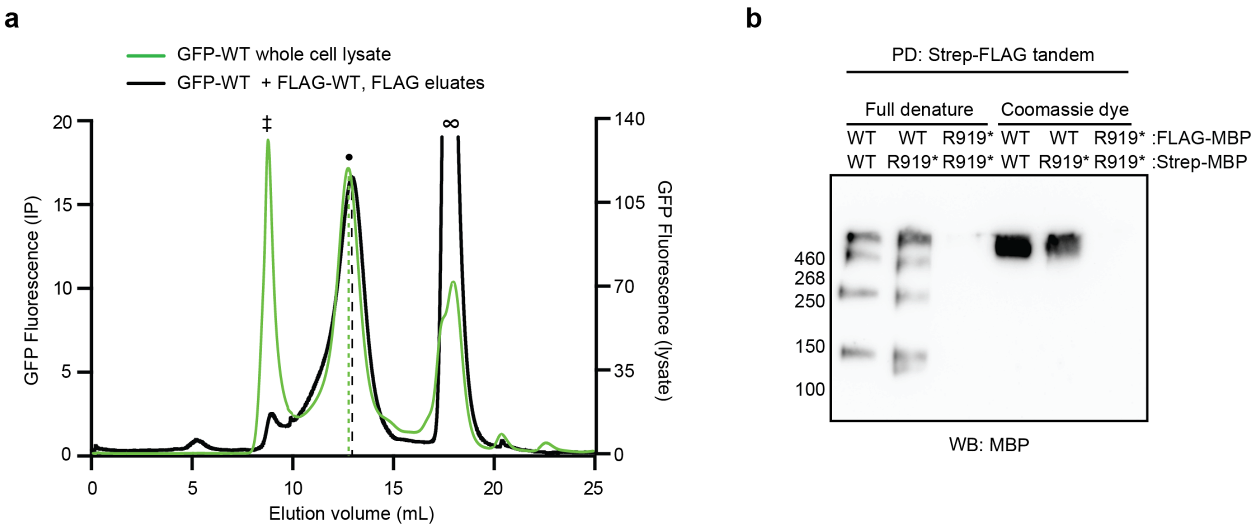


**Extended Data Figure 7. FSEC analysis WT TRPA1 complexes and Native Blue PAGE of WT and WT-R919* TRPA1 complexes.** (a) FSEC chromatograms of whole cell lysate from HEK293T cells transiently transfected with GFP-WT hTRPA1 (green trace) or FLAG immunoprecipitated eluates from HEK293T cells transiently co-transfected with GFP-WT hTRPA1 and 3xFLAG-WT hTRPA1 (black trace). Peaks corresponding to void (‡), tetrameric WT hTRPA1 channels (•) and free GFP (∞) are indicated. Dashed lines denote the center elution volume of each TRPA1 peak. (b) Immunoblotting analysis of tandem-purified WT/WT, WT/R919*, and R919*/R919* hTRPA1 complexes. The indicated FLAG-MBP-tagged and Strep-MBP-tagged constructs were transiently transfected in HEK293T cells. Lysates were tandem purified for Strep- then FLAG-tagged proteins. Eluents were run on a Blue Native PAGE gel after full (left) or partial (right) denaturation. MBP-tagged proteins of tandem purification eluents were probed using anti-MBP antibody. Data representative of 2 independent experiments.

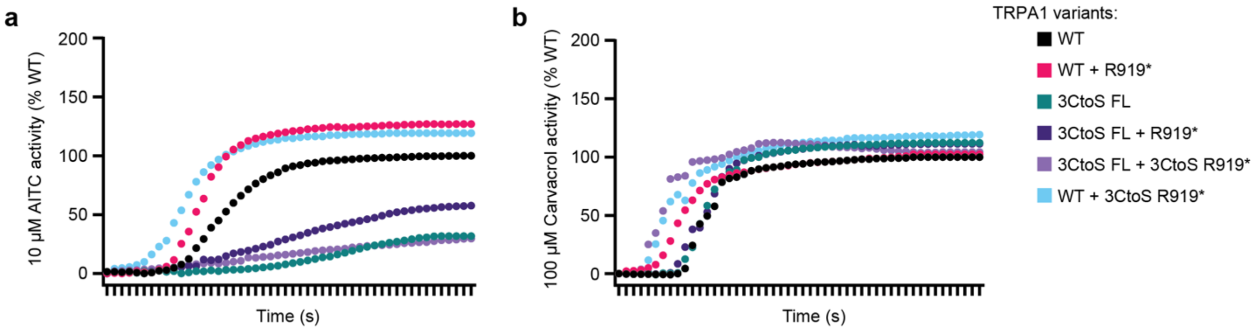


**Extended Data Figure 8. R919* TRPA1 subunits directly contribute to functional channels.** (a-b) Ratiometric calcium imaging traces of HEK293T cells transiently transfected with empty vector (mock), WT hTRPA1, 3CtoS FL hTRPA1, WT and R919* hTRPA1, 3CtoS FL and R919* hTRPA1, 3CtoS FL and 3CtoS R919* hTRPA1, or WT and 3CtoS R919* hTRPA1 from data quantified in Figure 5b. Cells were stimulated with 10 µM AITC (a) or 100 µM Carvacrol (b). Traces are averages of one representative experiment (n ≥ 60 cells) per condition. Data further normalized to WT hTRPA1 response.

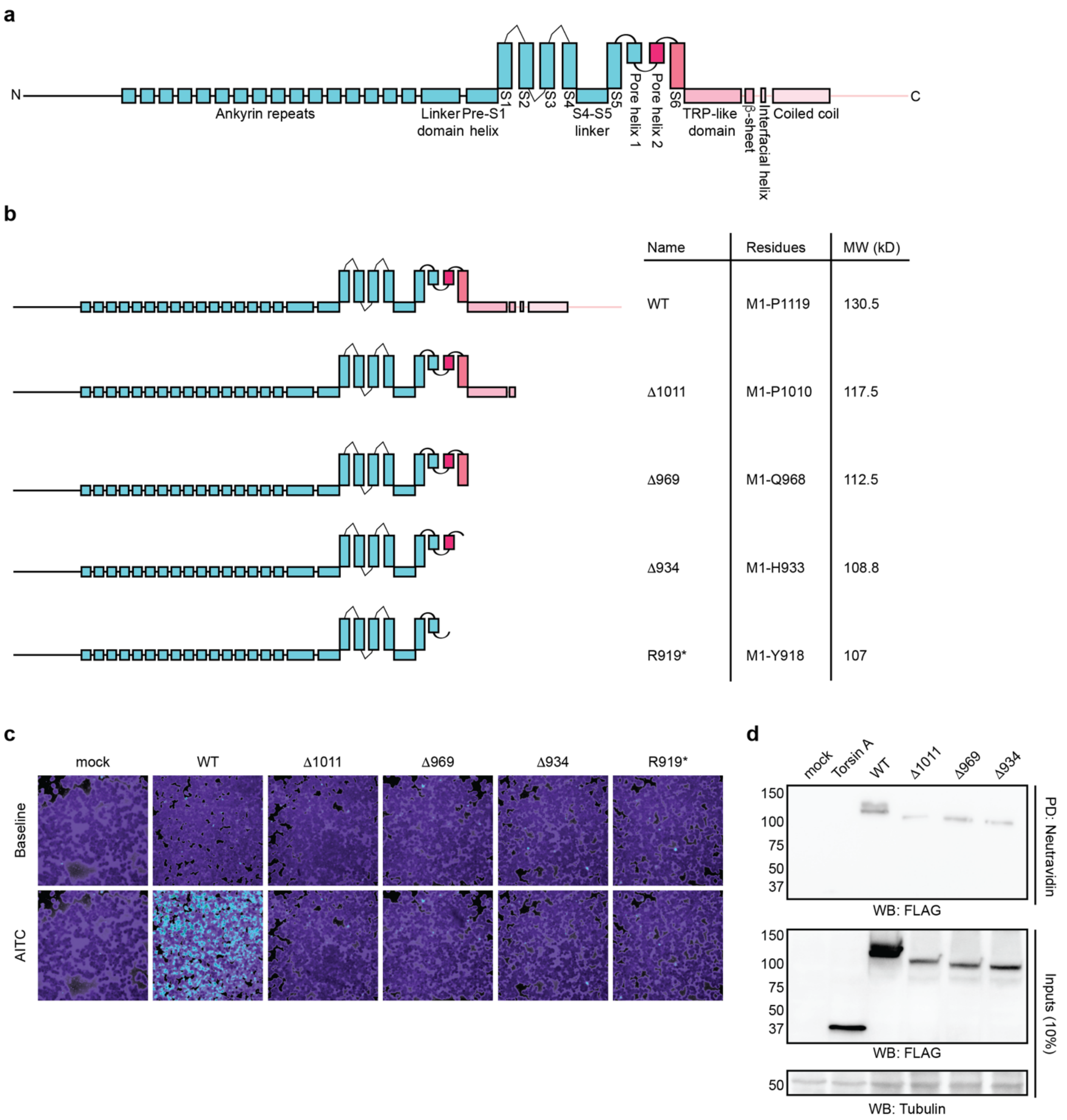


**Extended Data Figure 9. Design, activity, expression, and surface localization of C-terminal TRPA1 truncation constructs.** (a) Linear diagram depicting major structural domains in a WT hTRPA1 monomer. (b) Schematic representation and summary of composition of C-terminal TRPA1 truncation mutants assayed in Figure 6b and c. (c) Ratiometric calcium imaging of HEK293T cells transiently transfected with empty vector (mock) or the indicated 3xFLAG-tagged hTRPA1 constructs. Cells were stimulated with AITC (100 μM, n=60 cells per transfection condition). (d) Immunoblotting analysis of 3xFLAG-tagged hTRPA1 constructs or FLAG-Torsin A protein expression in biotin-labeled plasma membranes from transiently transfected HEK293T cells. Biotinylated proteins were precipitated by Neutravidin resin pulldown and probed using HRP-conjugated anti-FLAG antibody. Tubulin from whole cell lysates (10%, inputs) was the loading control. Torsin A was the negative control for plasma membrane localization. Data is representative of three independent experiments.

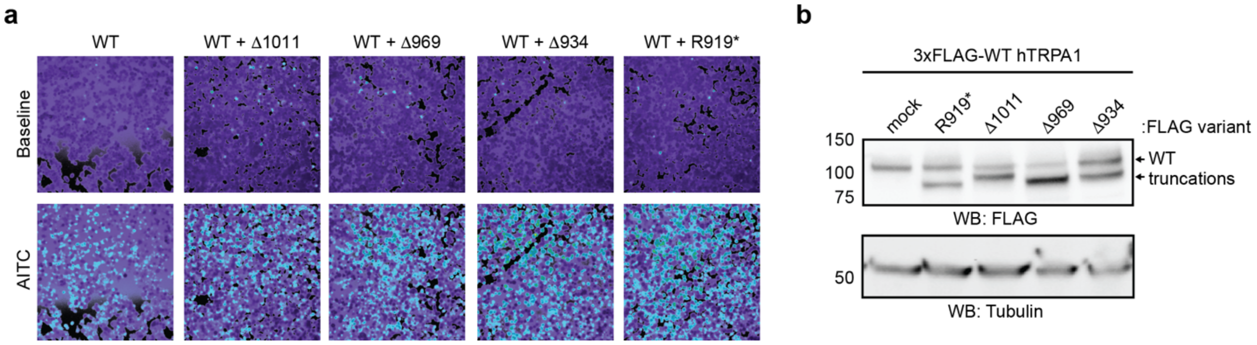


**Extended Data Figure 10. Mechanistic dissection of CFS-associated TRPA1 mutant-conferred channel hyperactivity.** (a) Ratiometric calcium imaging of HEK293T cells transiently co-transfected with 3xFLAG-WT hTRPA1 and empty vector (mock) or the indicated C-terminal truncation constructs from data quantified in Figure 6c. Cells were stimulated with 10 μM AITC. n ≥ 90 cells per condition. (b) Western blot of lysates from transiently transfected HEK293T cells expressing 3xFLAG-tagged hTRPA1 variants from (A), probed using HRP-conjugated anti-FLAG antibody. Tubulin was the loading control.

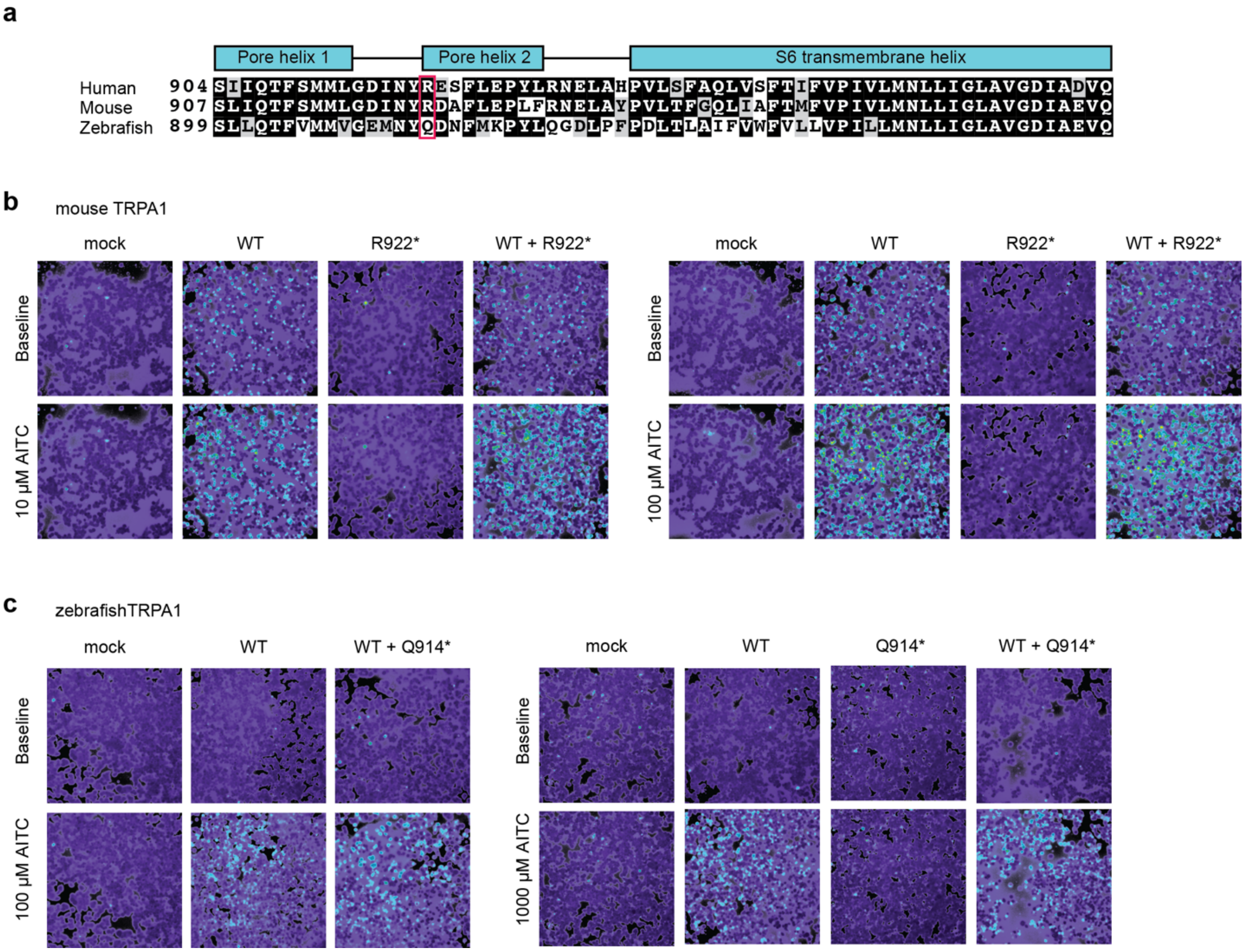


**Extended Data Figure 11. Evolutionary conservation of CFS-associated TRPA1 mutant-conferred channel hyperactivity in TRPA1 species orthologues.** (a) Alignment of mouse TRPA1 and zebrafish TRPA1a isoform with the human TRPA1 protein with protein topology indicated above. Amino acid residues are highlighted in black when present in the three proteins and in gray when they are present in only two sequences. The human R919 residue is indicated with a pink box. Alignment was built with T-Coffee [81] and BOXSHADE. (b) Ratiometric calcium imaging of HEK293T cells transiently transfected with empty vector (mock), WT mTRPA1, R922* mTRPA1, or WT and R922* mTRPA1 from data quantified in Figure 6d. Cells were stimulated with 10 µM (left) or 100 µM (right) AITC. n ≥ 90 cells per concentration. (c) Ratiometric calcium imaging of HEK293T cells transiently transfected with empty vector (mock), WT zTRPA1a, Q914* zTRPA1a, or WT and Q914* zTRPA1a from data quantified in Figure 6e. Cells were stimulated with 100 µM (left) or 1000 µM (right) AITC. n ≥ 90 cells per concentration.

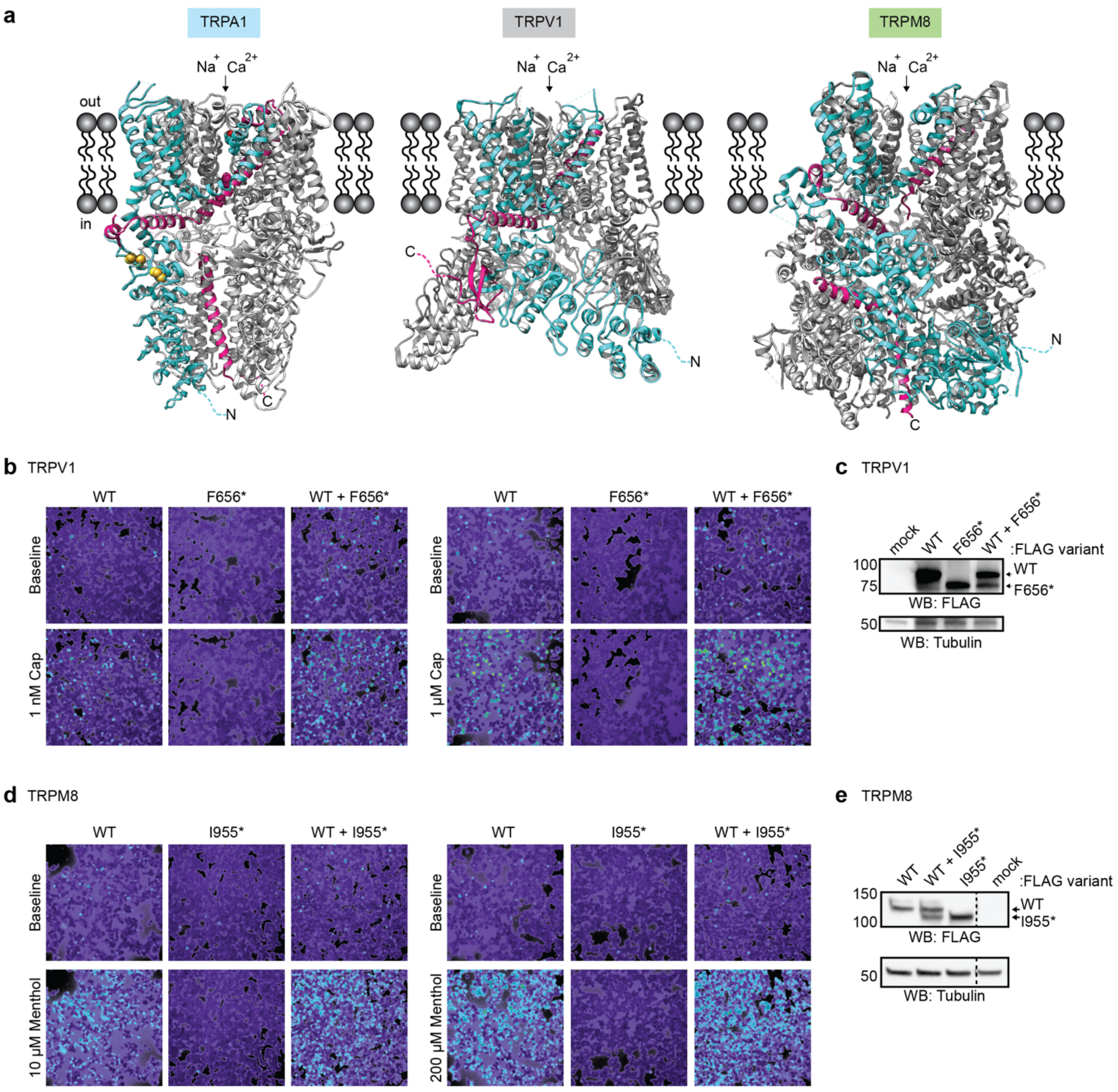


**Extended Data Figure 12. A TRPV1 mutant, but not a TRPM8 mutant, lacking the S6 transmembrane helix and cytoplasmic C-terminus confers channel hyperactivity with WT protein.** (a) Ribbon diagrams of TRPA1, TRPV1, and TRPM8. Regions retained in R919* hTRPA1, and the S6 and cytoplasmic C-terminus truncations for TRPV1 and TRPM8 are indicated in teal. Regions truncated in these mutants are indicated in pink. Only one subunit is colored for clarity. Models built with the human TRPA1 (PDB: 6V9W), rat TRPV1 (PDB: 7LP9), and *Parus major* TRPM8 (6O6A) Cryo-EM structures in UCSF Chimera. (b) Ratiometric calcium imaging of HEK293T cells transiently transfected with empty vector (mock), 3xFLAG-WT human TRPV1, 3xFLAG-F656* human TRPV1, or 3xFLAG-WT and F656* human TRPV1 from data quantified in Figure 6f. Cells were stimulated with 1 nM (left) or 1 µM (right) Capsaicin. n ≥ 90 cells per condition. (c) Western blot of lysates from transiently transfected HEK293T cells from (a), probed using HRP-conjugated anti-FLAG antibody. Tubulin was the loading control. (d) Ratiometric calcium imaging of HEK293T cells transiently transfected with empty vector (mock), 3xFLAG-WT rat TRPM8, 3xFLAG-I955* rat TRPM8, or 3xFLAG-WT and I955* rat TRPM8 from data quantified in Fig. 6g. Cells were stimulated with 10 µM (left) or 200 µM (right) Menthol. n ≥ 90 cells per condition. (e) Western blot of lysates from transiently transfected HEK293T cells from (c), probed using HRP-conjugated anti-FLAG antibody. Tubulin was the loading control.

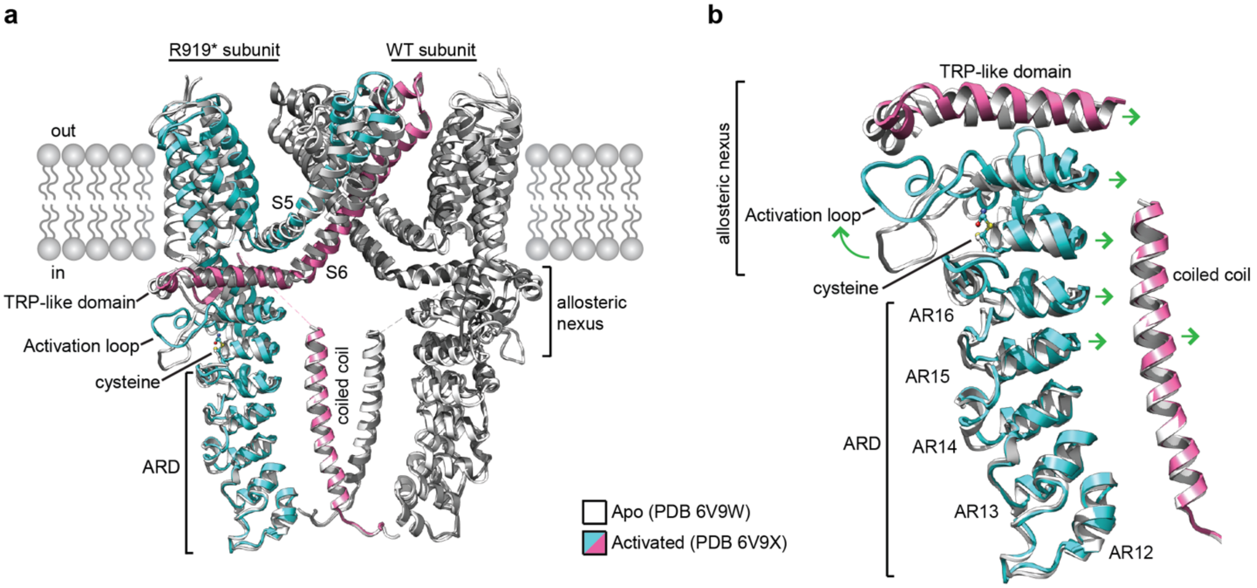


**Extended Data Figure 13. Conformational changes associated with TRPA1 activation.** (a) Overlay of ribbon diagrams of opposing R919* or WT TRPA1 in the closed (Apo, PDB 6V9W, white) and activated (PDB 6V9X, blue/pink or dark gray) states, respectively. Regions retained in R919* hTRPA1 are indicated in teal. Regions truncated in this mutant are indicated in pink. Only two opposing subunits are shown for clarity. Models built with the indicated TRPA1 Cryo-EM structures in UCSF Chimera. (b) Overlay of ribbon diagrams of the TRPA1 allosteric nexus, membrane-proximal ankyrin repeat domain (ARD), and coiled coil colored as in (a). Green arrows indicate regions and direction of gating associated conformational changes.
